# Beta cells are essential drivers of pancreatic ductal adenocarcinoma development

**DOI:** 10.1101/2024.11.29.626079

**Authors:** Cathy C. Garcia, Aarthi Venkat, Daniel C. McQuaid, Sherry Agabiti, Alex Tong, Rebecca L. Cardone, Rebecca Starble, Akin Sogunro, Jeremy B. Jacox, Christian F. Ruiz, Richard G. Kibbey, Smita Krishnaswamy, Mandar Deepak Muzumdar

## Abstract

Pancreatic endocrine-exocrine crosstalk plays a key role in normal physiology and disease. For instance, endocrine islet beta (β) cell secretion of insulin or cholecystokinin (CCK) promotes progression of pancreatic adenocarcinoma (PDAC), an exocrine cell-derived tumor. However, the cellular and molecular mechanisms that govern endocrine-exocrine signaling in tumorigenesis remain incompletely understood. We find that β cell ablation impedes PDAC development in mice, arguing that the endocrine pancreas is critical for exocrine tumorigenesis. Conversely, obesity induces β cell hormone dysregulation, alters CCK-dependent peri-islet exocrine cell transcriptional states, and enhances islet proximal tumor formation. Single-cell RNA-sequencing, *in silico* latent-space archetypal and trajectory analysis, and genetic lineage tracing *in vivo* reveal that obesity stimulates postnatal immature β cell expansion and adaptation towards a pro-tumorigenic CCK+ state via JNK/cJun stress-responsive signaling. These results define endocrine-exocrine signaling as a driver of PDAC development and uncover new avenues to target the endocrine pancreas to subvert exocrine tumorigenesis.

## INTRODUCTION

The pancreas is comprised of two functionally distinct cellular compartments: 1) the *endocrine* pancreas, composed of islet cells that secrete hormones to maintain glucose homeostasis, and 2) the *exocrine* pancreas, encompassing acinar cells that produce digestive enzymes and the ducts through which these enzymes traverse. Recent studies have challenged the longstanding dogma that the endocrine and exocrine compartments function independently, demonstrating endocrine-exocrine crosstalk in normal physiology and disease^1,2^. For example, maturity-onset diabetes of the young type 8 (MODY 8) is associated with mutations of *CEL*, a gene that encodes for carboxyl ester lipase produced and secreted by exocrine acinar cells^3^. Mutant CEL is acquired by islet beta (β) cells and aggregates intracellularly to induce endoplasmic reticulum (ER) stress, resulting in β cell dysfunction and diabetes, a disease of the endocrine pancreas^4^. Other exocrine diseases have similarly been associated with the development of diabetes, including chronic pancreatitis, Wolcott-Rallison syndrome, cystic fibrosis, and pancreatic ductal adenocarcinoma (PDAC)^2^. Conversely, obesity and diabetes – host metabolic states linked to β cell dysfunction – are associated with an increased risk of developing and dying of PDAC^5–10^, a highly lethal tumor primarily arising from acinar cells of the exocrine pancreas^11,12^. However, the cellular and molecular mechanisms that govern endocrine-exocrine signaling in tumorigenesis are not completely understood.

To date, few studies have directly examined the importance of β cells and β cell-secreted hormones in PDAC pathogenesis. Recent experiments have demonstrated a functional role of basal insulin in pancreatic tumorigenesis. Partial knockout of the insulin genes (*Ins1* and *Ins2*) or acinar-specific knockout of the insulin receptor (*InsR*) reduces oncogenic *Kras*-driven tumor development in mice fed a high-fed diet (HFD)^13–15^. Mechanistically, insulin signaling in acinar cells increases digestive enzyme production and acinar-to-ductal metaplasia (ADM)^15^, an early prerequisite step in PDAC development^12,16^. Our lab previously showed that obesity (due to loss of the appetite suppression hormone leptin (*Lep^ob/ob^*)^17^) enhances tumor progression and induces aberrant β cell expression of the peptide hormone cholecystokinin (CCK) in *Pdx1-Cre; Kras^LSL-G12D^; Lep^ob/ob^* (*KCO*) mice. Like insulin, CCK stimulates acinar cell proliferation, digestive enzyme production, and ADM^18–20^, and exogenous administration of the CCK analogue cerulein promotes oncogenic *Kras*-driven tumorigenesis in mice^21^. In *KCO* mice, β cell CCK expression – rather than insulin – strongly correlated with PDAC progression. Furthermore, induced weight loss or treatment with the antidiabetic dapagliflozin reduced β cell CCK expression and tumor formation^18^. Finally, transgenic β cell overexpression of CCK in lean *Pdx1-Cre; Kras^LSL-G12D^* (*KC*) mice was sufficient to increase acinar cell proliferation and ductal tumorigenesis^18^. These data argue that β cell-derived CCK is both a marker and driver of enhanced exocrine tumorigenesis.

In this study, we extend this work and bring new insights into the cellular and molecular pathways that regulate endocrine-exocrine signaling in PDAC progression. Specifically, we demonstrate that β cells play a critical role in PDAC development, as β cell ablation impedes exocrine tumorigenesis. Leveraging single-cell RNA sequencing (scRNA-seq) of multiple congenic obesity models and a suite of new machine learning-based computational tools for *in silico* lineage tracing (TrajectoryNet)^22,23^, latent-space archetypal analysis (AAnet)^24,25^, batch integration across datasets (scMMGAN)^26^, and optimal-transport distance-preserving embedding of patient data (DiffusionEMD)^27^, we gain deep insights into β and acinar cell heterogeneity, the dynamic transcriptional changes that lead to a pro-tumorigenic CCK+ β cell phenotype in obesity, and the resultant alterations in acinar cell states (**Figure S1**). Through both *in silico* (TrajectoryNet and AAnet) and experimental lineage tracing approaches and gene regulatory analysis, we further show that the CCK+ β cell state emerges from stress-induced expansion and adaptation of a postnatal immature β cell population. scMMGAN-enabled batch integration and DiffusionEMD-based patient embedding demonstrate that the obesity trajectory aligns with scRNA-seq datasets from both physiologic and pharmacologic β cell stressors across species. Finally, we show that pro-tumorigenic β cell CCK expression is mediated by stress-responsive JNK/cJun signaling via a novel distal *CCK* enhancer. Together, our results establish the critical importance of β cells in PDAC pathogenesis and implicate β cell stress pathways that could be targeted to intercept exocrine tumorigenesis.

## RESULTS

### β cell loss suppresses pancreatic exocrine tumorigenesis

β cells are ideally positioned within pancreatic islets to signal to neighboring exocrine acinar cells to drive PDAC development, yet whether β cells are required for exocrine tumorigenesis has not been firmly established. To test this, we crossed the *KPC* (*Pdx1-Cre; Kras^LSL-G12D^; Trp53^LSL-R^*^172^*^H/+^*) mouse model that faithfully recapitulates the genetic and histologic progression of human PDAC^28^ with the *Akita* model (**Figure 1A**). *Akita* mice harbor a mutation in the *Ins2* (insulin) gene, which causes protein misfolding, ER stress, and, consequently, non-inflammatory β cell death^29^. We confirmed decreased β cell mass and induction of ER stress in islets of *KPC-Akita* mice (**Figure 1B**). Strikingly, *KPC-Akita* mice exhibited reduced overall tumor burden compared to age-matched *KPC* littermates (**Figure 1C**). Despite decreased tumorigenesis, *KPC-Akita* mice showed markedly elevated glucose levels (**Figure 1D**), suggesting that hyperglycemia is not independently pro-tumorigenic in the absence of β cells. These data showcase an essential role for β cells in PDAC progression.

**Figure 1.**
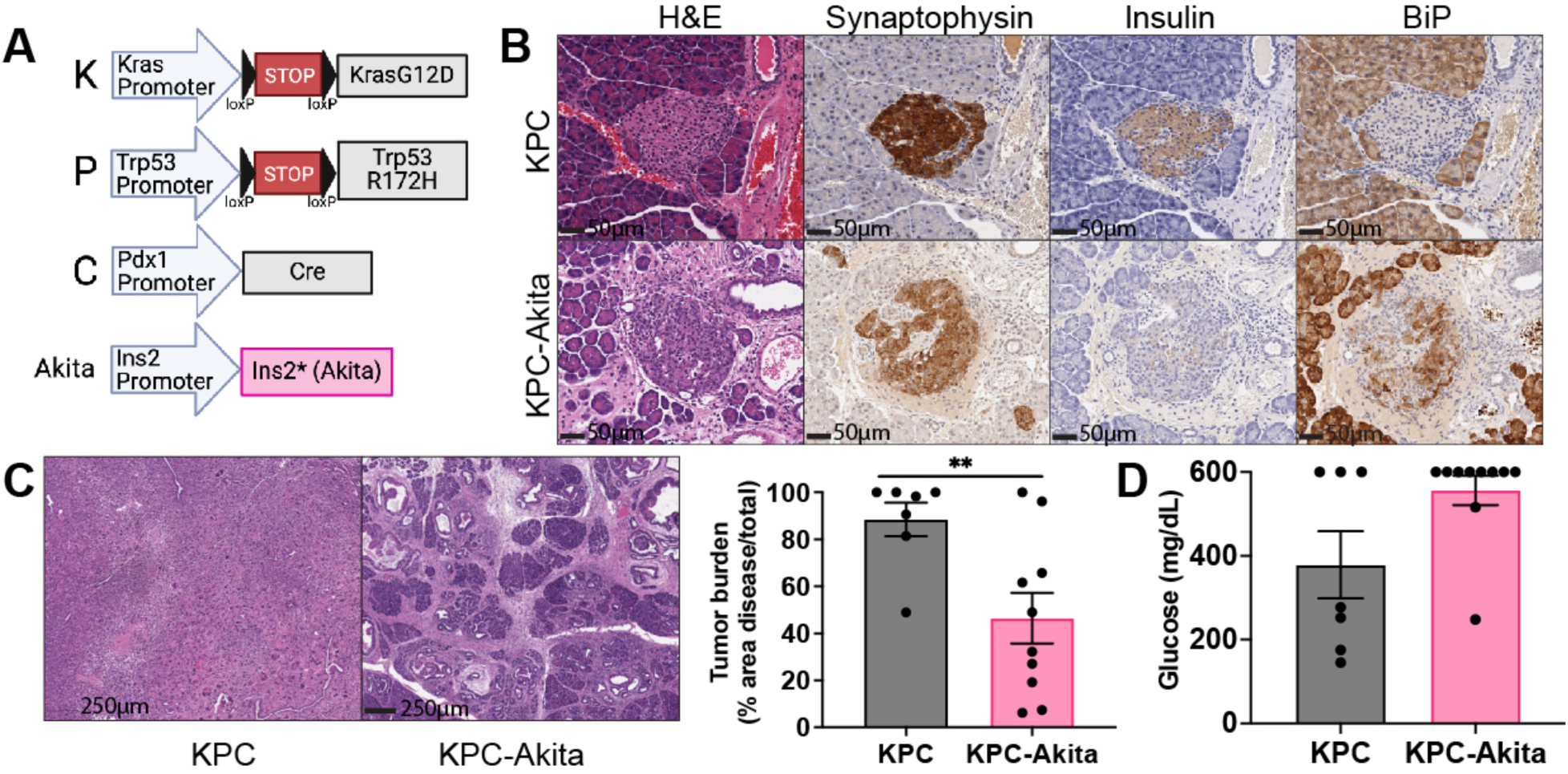
Loss of β cells impedes pancreatic tumorigenesis. (A) Schematic of the alleles used to generate *KPC* and *KPC-Akita* mice (generated with Biorender). (B) H&E and IHC stains show loss of insulin-expressing β cells and increased ER stress (BiP) in islets (labeled by pan-neuroendocrine marker synaptophysin) of *KPC-Akita* mice. (C) Representative H&E images of *KPC* and *KPC-Akita* mice and quantification of tumor burden (mean +/- s.e.m., n=7-10) at 12 weeks of age. \*\**p*<0.01, Welch’s t test. (D) Final random blood glucose (mean +/- s.e.m., n=7-10, maximum glucometer measurement is 600 mg/dL) of mice, no significance, Welch’s t test.

### Dynamic changes in β cell expression of insulin and CCK with obesity

Having established that β cells are basally required for exocrine tumorigenesis, we next investigated how obesity, a pro-tumorigenic host metabolic state^18,30–34^, modulates β cells to drive tumorigenesis. We performed scRNA-seq of isolated islet cells from aged-matched congenic (C57/B6) male wild-type (WT), moderately obese HFD-fed, and severely obese *Lep^ob/ob^* mice that displayed increasing weight and glucose levels (**Figures S2A-S2C**). Consistent with these phenotypic observations, primary islets isolated from HFD-fed and *Lep^ob/ob^* mice exhibited decreased glucose-stimulated insulin secretion (GSIS), a marker of β cell dysfunction (**Figures S2D** and **S2E**). After pre-processing of scRNA-seq data (see **Methods**), we visualized and clustered 23,469 single cells into 20 clusters (**Figure S2F**). We annotated these clusters based on published endocrine and exocrine markers^35^ (**Figure S2G**) and subsetted the data to 20,294 endocrine cells for downstream analysis.

We re-embedded and clustered the endocrine cells, annotating clusters corresponding to alpha (ɑ), β, delta (δ), and PP cells based on high expression of glucagon (*Gcg*), insulin (*Ins1, Ins2*), somatostatin (*Sst*), and polypeptide (*Ppy*), respectively (**Figure S3A**). Similar to prior studies^36,37^, a small population of polyhormonal cells (expressing insulin and a second hormone) was also observed. CCK (mouse gene *Cck*) was primarily expressed in β cells, the predominant cell type of the endocrine pancreas, and β cells showed the greatest variation in transcriptional states across the conditions (**Figure S3A**). Given this, we re-embedded β cells from the three conditions (**Figure S3B**). Then, we continuously inferred individual and heterogeneous cellular trajectories over the progression of obesity using an ODE-based dynamic optimal transport neural network termed TrajectoryNet^22,23^. TrajectoryNet is designed to interpolate continuous dynamics for every single cell from sampled timepoints, where the dynamics learned are biologically plausible through ensuring energy efficiency and modeling cellular proliferation (see **Methods**). This represents an advancement over traditional trajectory-based inference methods, which calculate a single pseudotemporal ordering over the entire population of cells^38^. Here, we learned individual cellular trajectories from WT (timepoint 1) to HFD (timepoint 2) to *Lep^ob/ob^* (timepoint 3) conditions. Then, the transcriptional changes along the learned trajectories were analyzed by decoding them back into the gene expression space. This analysis revealed a clear trajectory corresponding to the obesity progression axis, wherein *Ins1* and *Ins2* showed decreasing expression while *Cck* showed increasing expression along the trajectory (**Figure S3B**). Gene trends learned by TrajectoryNet showed a decrease in β cell identity, maturation, and insulin secretion markers and increased protein processing, dedifferentiation, and ER stress markers as obesity progresses (**Figure S3C**). These data argue that β cells retain secretory capacity but upregulate *Cck* at the expense of insulin. We took advantage of the Min6 murine insulinoma cell line to directly detect hormones within secretory granules. This frequently employed β cell model expresses insulin and CCK, comparable to β cells of tumor-bearing *KCO* mice (**Figure S3D**). Co-immunoelectron microscopy revealed that insulin and CCK were present in the same granules, including those at the plasma membrane (**Figure S3E**). These findings indicate that these hormones may be co-secreted by β cells with the capacity to signal to neighboring exocrine cells.

### Peri-islet acinar cell adaptation and transformation in obese mice

To determine whether these β cell adaptations to obesity correlate with transcriptional changes in exocrine acinar cells, the putative cell-of-origin for PDAC^11,12,16^, we re-embedded scRNA-seq data for *Cpa1+/Prss2+* cells. We recovered ∼250 acinar cells that passed quality control metrics from all samples (**Figure 2A**). As these cells were obtained from islet isolations and represented ∼1% of sequenced cells, we reasoned that they likely surrounded the islet (peri-islet). Consistent with this hypothesis, acinar cells from obese (HFD-fed and *Lep^ob/ob^*) versus lean (WT) mice were enriched in gene signatures observed in peri-islet acinar cells derived from the leptin receptor-deficient (*Lepr^db/db^*) obesity model^39,40^ (**Figure 2B**). Differential expression analysis revealed multiple proteases with increased expression in acinar cells of HFD-fed and *Lep^ob/ob^* mice (**Figure 2C**). These included proteases induced by CCK signaling in acinar cell models (*Cela3b, Cpb1, Ctrb1*) and spatially confirmed to be upregulated in peri-islet acinar cells (*Try4*, *Try5*, *Try10*) in obese *Lepr^db/db^* mice by single molecule fluorescence *in situ* hybridization (smFISH)^40^.

**Figure 2.**
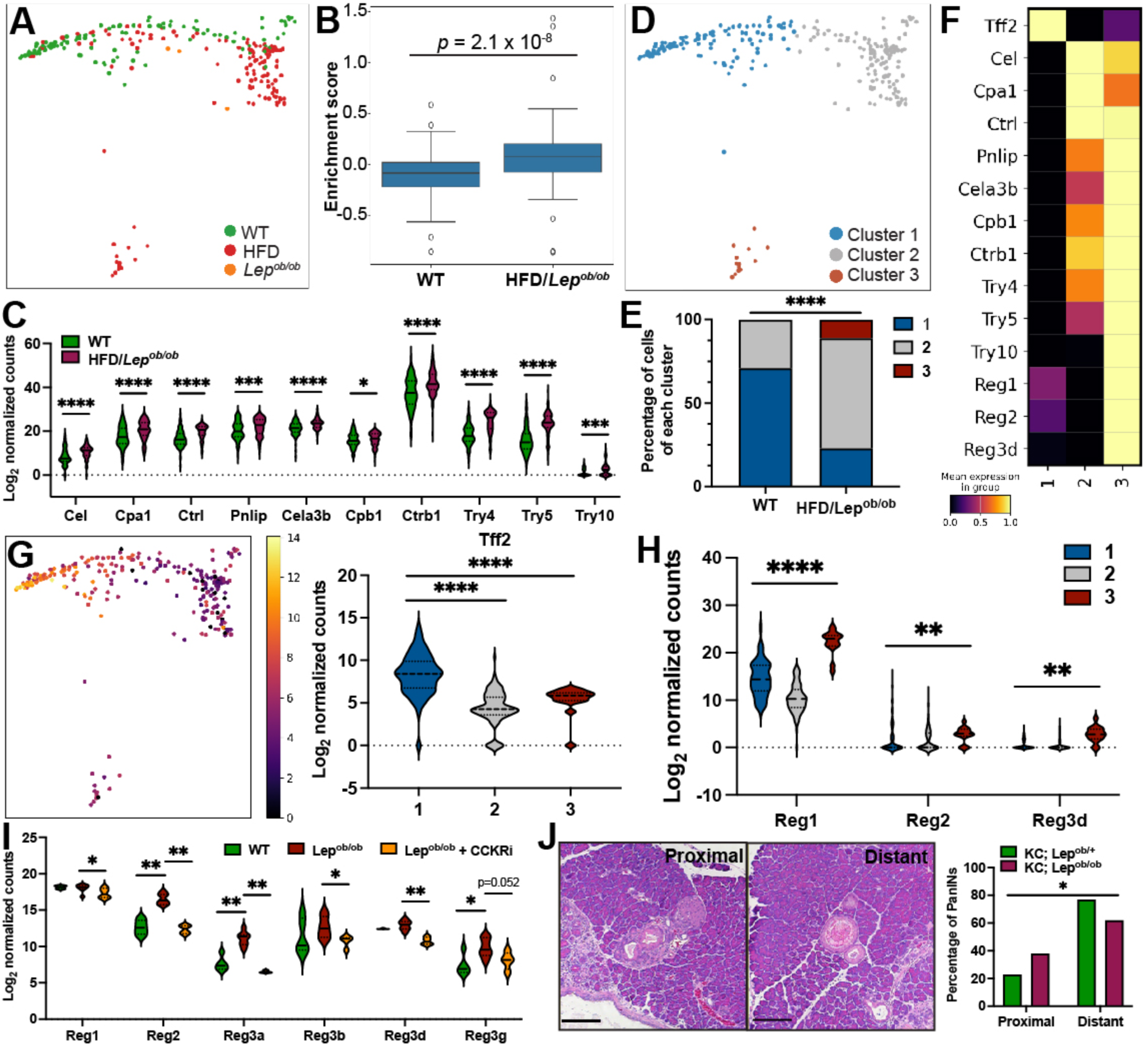
Peri-islet acinar cell adaptation and transformation in obesity. (A) PHATE visualization of *Cpa1+/Prss2+* acinar cells coded by sample condition. (B) Enrichment of gene signatures observed in bulk RNA-seq of peri-islet acinar cells from *Lepr^db/db^* mice (log_2_FC expression>1.5, FDR<0.05)^40^ in obesity models. Box plots display 25^th^, 50^th^, and 75^th^ percentile enrichment scores +/- 1.5 interquartile range (IQR) for each cell in each condition. *p*-value is derived from one-sided Wilcoxon rank sum test. (C) Proteases are upregulated in HFD/*Lep^ob/ob^* (n=140 cells) versus WT (n=112 cells) acinar cells. Violin plots represent gene expression distribution (min/max with 25^th^, 50^th^, and 75^th^ percentiles delineated by lines). \**q*<0.05, \*\*\**q*<0.001, \*\*\*\**q*<0.0001, Wilcoxon rank sum test adjusted for multiple comparisons with two-stage Benjamini, Krieger, Yekutieli step-up. Notably, *Cela3b*, *Cpb1*, and *Ctrb1* are induced by CCK in AR42J acinar cell carcinoma cells and *Try4*, *Try5*, and *Try10* were shown to enriched in peri-islet acinar cells in *Lepr^db/db^* mice^40^. (D) PHATE visualization of *k*-means clustering of acinar cells in (**A**) derived from lean (WT) vs. obese (HFD/*Lep^ob/ob^*) conditions. (E) Differential proportions of acinar cell clusters in (**D**) ****p<0.0001, chi-square test. (F) Heatmap of row normalized mean expression of *Tff2*, proteases, and *Reg* genes across cells within each cluster in (**D**). (G) PHATE visualization and violin plots of *Tff2* expression in acinar cell clusters. \*\*\*\**p*<0.0001, Wilcoxon rank sum test. (H) *Reg* gene expression in acinar cells in (**D**). Violin plots represent gene expression distribution (min/max with 25^th^, 50^th^, and 75^th^ percentiles delineated by lines). \*\**q*<0.01, \*\*\*\**q*<0.0001, Wilcoxon rank sum test adjusted for multiple comparisons with two-stage Benjamini, Krieger, Yekutieli step-up. (I) Violin plots of pancreatic *Reg* gene expression by RNA-seq of 12-week-old C57/B6 WT (n=5), *Lep^ob/ob^* (n=7), and *Lep^ob/ob^* mice treated with CCK receptor antagonists (proglumide, lorglumide; n=2 mice/drug) for 6 weeks. \**q*<0.05, \*\**q*<0.01. Wilcoxon rank sum test adjusted for multiple comparisons with two-stage Benjamini, Krieger, Yekutieli step-up. (J) Representative images and percentage of islet proximal (<400 µm) or distant (>400 µm) PanINs in lean 13-week-old *KC; Lep^ob/+^*and obese 7-week-old *KC; Lep^ob/ob^* mice (n=4-5 mice/group). \**p*<0.05, two-sided Fisher’s exact test. Scale bars are 100 µm.

We next clustered acinar cells into three subclusters, the proportions of which were altered by obesity (**Figures 2D** and **2E**). Obesity was associated with a decrease in *Tff2*+ acinar cells (cluster 1) (**Figures 2F** and **2G**), which have been shown to be resistant to oncogenic *Kras*-induced transformation^41^. Conversely, obesity was associated with an increase in the proportion of acinar cells expressing higher levels of proteases (clusters 2 and 3) (**Figures 2F**). Strikingly, cluster 3 was uniquely present in obesity models (**Figure 2E**), and cells within this cluster exhibited high expression levels of the Regenerating (*Reg*) gene family (**Figures 2F** and **2H**). *Reg* expression also occurs in peri-islet acinar cells in human pancreas samples^42^ and has been associated with acinar cell stress and inflammation^43^. Additionally, Reg proteins can promote ADM^44,45^, making them both markers and drivers of a tumor-permissive acinar cell state. In concordance with our scRNA-seq data, bulk pancreata of obese *Lep^ob/ob^* mice exhibited upregulation of *Reg* genes compared to lean congenic WT mice (**Figure 2I**). Importantly, *Reg* expression was abolished by CCK receptor antagonist treatment of *Lep^ob/ob^* mice (**Figure 2I**), indicating that cluster 3 cells may be induced by signaling from β cell-derived CCK in mice. We hypothesized that these CCK-dependent alterations in peri-islet transcriptional states may enhance *Kras*-driven tumorigenesis in proximity to islets in obesity. As predicted, we found that early tumors in obese *KCO* mice emerged in greater proportions in proximity to islets than in lean *KC; Lep^ob/+^* mice (**Figure 2J**). These data support endocrine-exocrine CCK signaling in acinar cell adaptation and pancreatic tumor development.

### Identification of the cell-of-origin for pro-tumorigenic β cells in obesity

Given that *Cck*+ β cells correspond to a pro-tumorigenic phenotype^18^, we next sought to decipher how they emerge in response to obesity. Obesity increases insulin demand, augmenting β cell mass in mice and humans^46,47^. Lineage tracing studies in mice have shown that new β cells primarily arise from self-duplication rather than transdifferentiation from other endocrine or exocrine cell types under a variety of physiologic scenarios, including homeostasis, pregnancy, and pancreatic injury^48^. In contrast, upon complete β cell ablation, bi-hormonal endocrine cells emerge from α (*Gcg*+) and δ (*Sst*+) cells that acquire insulin expression and become mono-hormonal *Ins1/2*+ cells^49,50^, supporting islet cell plasticity in regeneration. Additional studies have demonstrated that exocrine duct cells can transdifferentiate into β cells upon pancreatic injury^51,52^. To distinguish whether *Cck*+ cells arise from β cell duplication versus transdifferentiation in obesity, we leveraged TrajectoryNet for *in silico* lineage tracing. Since TrajectoryNet outputs a trajectory for each cellular state, we can trace trajectories of pathogenic states backwards to find their cell-of-origin. We traced back the most likely paths of *Cck*+ *Lep^ob/ob^* cells, allowing us to identify “cell-of-origin” cells, defined as the cells at the first timepoint of the learned trajectories (**Figure 3A**, see **Methods**). We then compared the proportion of each cell type within the cells-of-origin versus the distribution in the measured WT cells, terming the ratio the cell type “enrichment score” (see **Methods**). This analysis revealed that *Cck*+ *Lep^ob/ob^* β cells derive from β cells from the WT condition, and *Cck*+ *Lep^ob/ob^* polyhormonal cells derive from polyhormonal cells from the WT condition (**Figure 3B**). Thus, *Cck*+ cells arise from pre-existing β cells rather than by transdifferentiation of other cell types.

**Figure 3.**
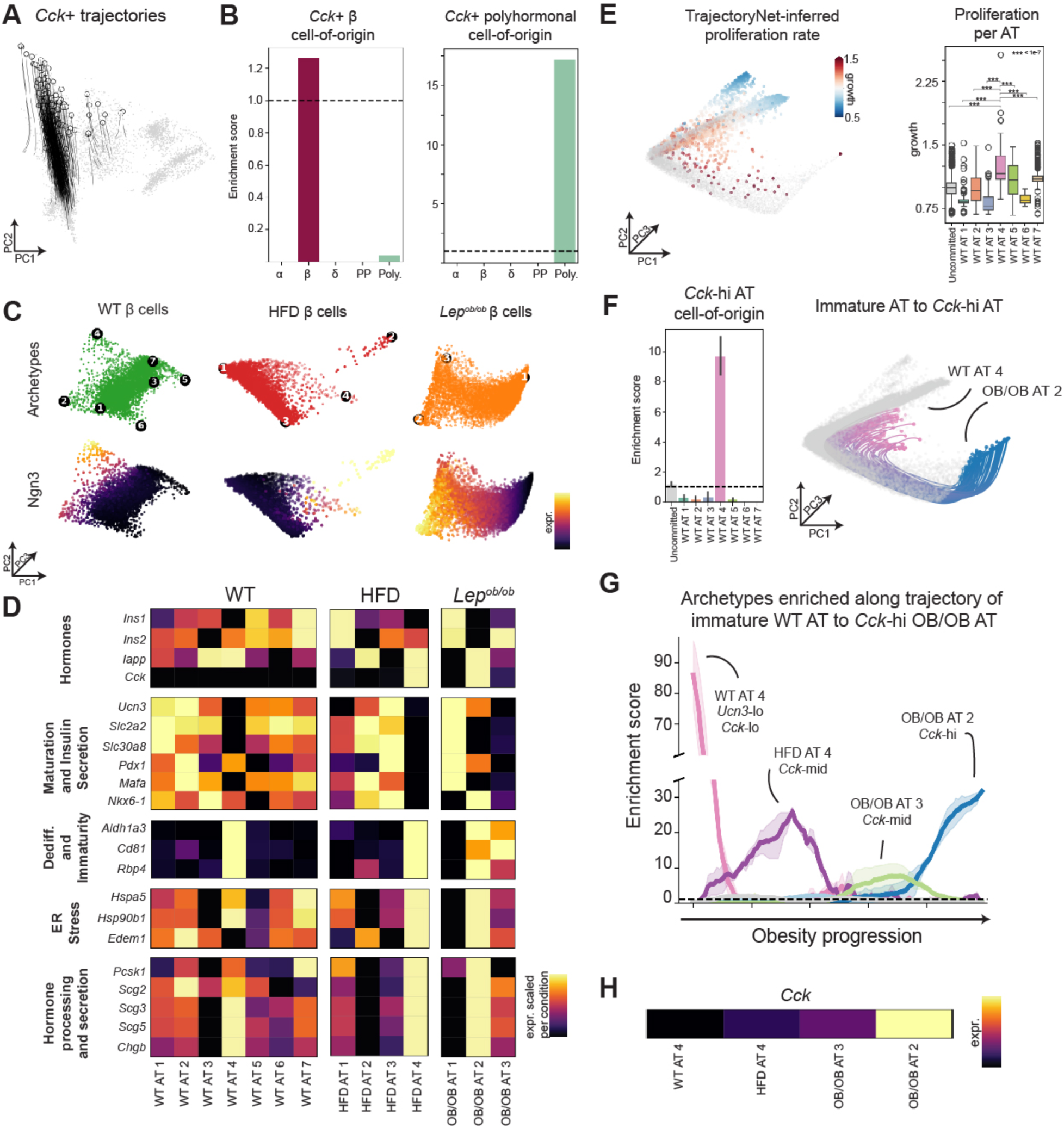
*In silico* identification of the cell-of-origin for pro-tumorigenic β cells in obesity. (A) TrajectoryNet-learned trajectories tracing paths from cells from WT mice to *Cck*+ *Lep^ob/ob^* cells. (B) Enrichment score of each islet cell type for cell-of-origin of *Cck*+ *Lep^ob/ob^* β cells (left) and *Cck*+ *Lep^ob/ob^* polyhormonal cells (right), calculated over two TrajectoryNet runs that yielded identical results. (C) AAnet-learned archetypes for WT, HFD, and *Lep^ob/ob^* samples (top row). Color embedding of denoised expression of Neurogenin 3 (*Ngn3*) across different archetypes. (D) Heatmap (row-normalized within each condition) for marker genes expression for each archetype in (**A**). (E) TrajectoryNet-inferred proliferation visualized for cells from WT condition, where proliferation rate > 1 implies growth. Cells committed to WT AT 4 have a significantly higher growth rate than other archetypes or uncommitted cells. Box plots display 25^th^, 50^th^, and 75^th^ percentiles +/- 1.5 interquartile range (IQR). \*\*\**p* < 1×10^-7^, Wilcoxon rank sum test. (F) Enrichment scores (mean +/- 95% confidence interval, n=2 TrajectoryNet runs) for each WT archetype within cells-of-origin for *Cck*-hi *Lep^ob/ob^* AT 2. High enrichment of WT AT 4 implies trajectories from WT AT 4 to *Lep^ob/ob^*AT 2 as shown. (G) Mean enrichment score for each archetype within cells on the trajectory from WT AT 4 to *Lep^ob/ob^*AT 2, calculated at each timepoint over two TrajectoryNet runs, demonstrating archetypes (HFD AT 4 and *Lep^ob/ob^* AT 3) that represent potential intermediate states. (H) Heatmap of row-normalized *Cck* expression for archetypes WT AT 4, HFD AT 4, *Lep^ob/ob^* AT 3, and *Lep^ob/ob^* AT 2.

To validate our *in silico* findings, we performed *in vivo* lineage tracing by crossing *Gcg-Cre^ERT^*^2^ and *Ins1-Cre^ERT^* mouse lines^53,54^ with *Lep^ob/ob^* mice harboring a Cre-inducible fluorescent reporter (*Rosa26^LSL-TdTomato^*)^55^ to label α and β cells, respectively (**Figures 4A, S4A,** and **S4B**). We treated mice with tamoxifen at ∼4 weeks of age (before significant obesity onset), aged all mice to 16 weeks, and performed co-immunofluorescence for CCK and insulin on isolated pancreatic sections to determine overlap with TdTomato+ cells (**Figures 4A** and **4B**). Importantly, all end glucose levels and weights were comparable across groups in *Lep^ob/ob^* mice (**Figures S4C and S4D**). *Gcg-Cre^ERT^*^2^ mice showed CCK expression in most of the islet, but only ∼3% of CCK+ cells overlapped with TdTomato+ cells (**Figures 4B** and **4C**), indicating that CCK+ cells were not primarily arising from ɑ to β cell transdifferentiation. In contrast, most CCK+ cells were labeled with TdTomato in *Ins1-Cre^ERT^* mice (**Figures 4B** and **4C**). The incomplete labeling of CCK+ cells in *Ins1-Cre^ERT^* mice was likely due to inefficient labeling of β cells with this *Cre^ERT^*line in obese mice (**Figure S4B**), which has been previously observed in lean mice^54^. In support of this hypothesis, we discovered a positive linear correlation between TdTomato+/insulin+ cells (a marker of labeling efficiency) and TdTomato+/CCK+ cells (**Figure 4D**). Collectively, these data argue that CCK+ cells arise from pre-existing β cells.

**Figure 4.**
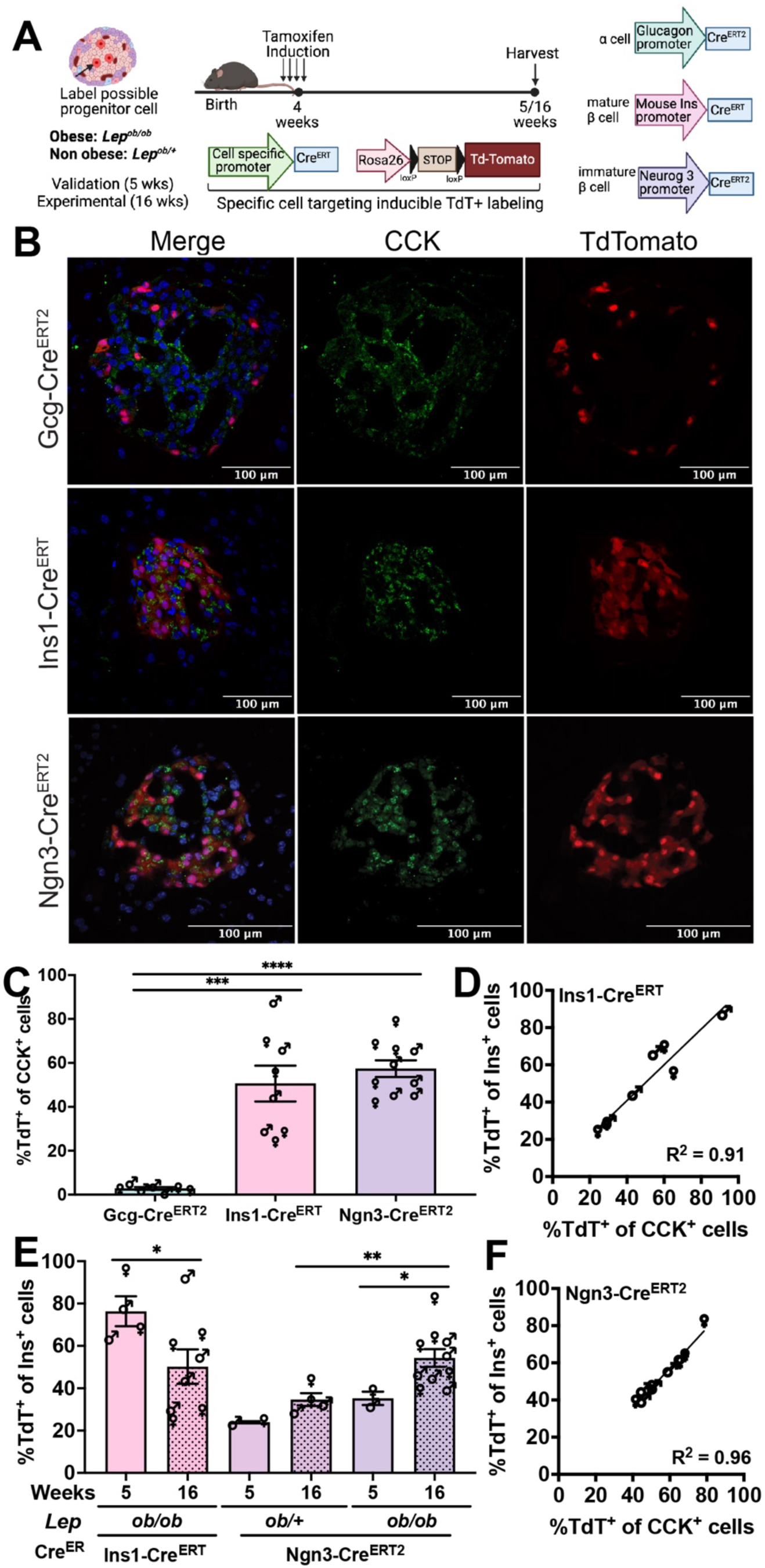
Immature β cells are the cell of origin for the pro-tumorigenic CCK+ β cell state. (A) Schematic of experimental setup and timeline of lineage tracing experiments (generated with Biorender.com). Mice were given tamoxifen at ∼4 weeks (prior to significant obesity) and analyzed at ∼5 weeks (one-week post-tamoxifen) to ensure fidelity of labeling or ∼16 weeks of age (when islet CCK expression is present) for lineage tracing analysis. (B) Representative images of 16-week-old *Gcg-Cre^ERT^*^2^*; Lep^ob/ob^; Rosa26^LSL-TdTomato^*, *Ins1-Cre^ERT^; Lep^ob/ob^; Rosa26^LSL-TdTomato^*, and *Ngn3-Cre^ERT^*^2^*; Lep^ob/ob^; Rosa26^LSL-TdTomato^* mice administered tamoxifen at 4 weeks of age and analyzed for CCK expression (by immunofluorescence) in lineage-traced TdTomato+ cells. DAPI labels nuclei blue. (C) Average percentage (mean+/-s.e.m., n=7-10 mice per group, sex is designated by symbols) of CCK+ cells labeled with TdTomato in mice in (B). ***p<0.001, ****p<0.0001, Welch’s t-test. (D) Strong positive correlation (Pearson R^2^: 0.91) between total labeled insulin+/TdTomato+ and CCK+/TdTomato+ cells in *Ins1-Cre^ERT^; Lep^ob/ob^; Rosa26^LSL-TdTomato^* mice in (**C**). (E) Average percentage (mean+/-s.e.m., n=2-10 mice per group, sex is designated by symbols) of insulin+ cells labelled with TdTomato in obese *Ins1-Cre^ERT^; Lep^ob/ob^; Rosa26^LSL-TdTomato^*, obese *Ngn3-Cre^ERT^*^2^*; Lep^ob/ob^; Rosa26^LSL-TdTomato^*, and lean *Ngn3-Cre^ERT^*^2^*; Lep^ob/+^; Rosa26^LSL-TdTomato^* mice analyzed at 5 (1 week post-tamoxifen) or 16 weeks (12 weeks post-tamoxifen). *p<0.05, **p<0.01, Welch’s t-test. (F) Strong positive correlation (Pearson R^2^: 0.96) between total labeled insulin+/TdTomato+ and CCK+/TdTomato+ cells in *Ngn3-Cre^ERT^*^2^*; Lep^ob/ob^; Rosa26^LSL-TdTomato^* mice in (**C**).

### A latent-space archetypal analysis network reveals β cell heterogeneity corresponding to maturation state and function

Recent studies have suggested that adult β cells are heterogeneous and that host metabolic conditions can modulate this heterogeneity^36,56,57^. To disentangle β cell heterogeneity with obesity, we separately re-embedded the cells for each condition (WT, HFD, and *Lep^ob/ob^*). We characterized heterogeneity with the latent-space archetypal analysis approach AAnet. AAnet is an autoencoder that embeds cells into a simplicial latent representation and defines them with respect to cellular “archetypes,” or extreme states^24,25^ (see **Methods**). Unlike traditional archetypal analysis, AAnet does not assume that cells are archetypal in gene expression space but instead finds a lower dimensional latent space that is amenable for such factor analysis. This enables the characterization of β cell states while preserving continuous variation between highly plastic ones. AAnet identified seven archetypes (AT) within cells from WT mice, four archetypes from HFD-fed mice, and three archetypes from *Lep^ob/ob^* mice, where the number of archetypes was determined based on the elbow point of reconstruction error^20^ (**Figures 3C** and **S5A**). Similarity scores calculated between all archetypes identified three groups with shared archetypal signatures (Groups 1, 2, and 3) and differing proportions across conditions (**Figure S5B** and **S5C)**.

We characterized these β cell archetypes using marker genes defined by previous research on β cell heterogeneity^56,58,59^ (**Figures 3D** and **S5D-S5F** and **Table S1**). Group 1 archetypes (WT ATs 5 and 7; HFD AT 1; *Lep^ob/ob^* AT 1) were significantly (*q* < 0.05) enriched for expression of insulin, insulin secretion markers, and β cell maturation markers, consistent with a mature β cell phenotype (**Figure 3D** and **Table S1**). Group 3 archetypes (WT ATs 1, 2, 3, and 6; HFD ATs 2 and 3) were characterized by significantly (*q* < 0.05) higher mitochondrial expression and – compared to Group 1 – reduced insulin expression (**Figure 3D** and **Table S1**). For WT AT 1 and HFD AT 3, maturation and insulin secretion markers *Ucn3*, *Slc2a2*, *Slc30a8* were significantly enriched. Group 2 archetypes (WT AT 4, HFD AT 4, *Lep^ob/ob^* AT 2, and *Lep^ob/ob^* AT 3) displayed significantly lower expression (versus other archetypes within each condition) of insulin (*Ins1*) and maturation markers and higher expression of dedifferentiation/immaturity^35,60^, hormone processing, and secretion markers (**Figure 3D** and **Table S1**). Together, these findings capture β cell diversity and complexity and reveal archetypal clusters of immature (Group 2), intermediate (Group 3), and mature (Group 1) β cells across WT, HFD, and *Lep^ob/ob^*mice.

### Immature β cells are the cell-of-origin for the pro-tumorigenic β cell state

Given significant β cell heterogeneity in these models, we next used learned cellular dynamics from TrajectoryNet to decipher which archetypes are most likely to be the cell-of-origin for *Cck-*hi *Lep^ob/ob^* cells (*Lep^ob/ob^* AT 2; **Figure 3D**). TrajectoryNet utilizes an auxiliary network to learn the relative proliferation rates of cells from the WT condition, where WT cells predicted to be proliferating have a relative growth rate of >1. Across all WT archetypes, cells committed to WT AT 4 had a significantly higher proliferation rate than all other archetypes (**Figure 3E**). This argues that immature β cells represent a regenerative subpopulation in adult mice, consistent with prior data^61,62^. We retrieved the cells-of-origin for the *Cck*-hi *Lep^ob/ob^* AT 2 cells and determined the enrichment for each WT archetype within this population (see **Methods**). Overwhelmingly, WT AT 4 was the most likely source for *Cck-*hi *Lep^ob/ob^* AT 2 cells over all other archetypes (**Figure 3F**), suggesting that immature β cells give rise to *Cck*-hi β cells in response to obesity. We next leveraged TrajectoryNet to derive intermediate states that may arise along this axis, computing the enrichment of archetypes along the trajectory of cells originating from WT AT 4 and terminating at *Lep^ob/ob^* AT 2 (see **Methods**). As expected, WT AT 4 and *Lep^ob/ob^* AT 2 were highly represented at the beginning and end of the trajectory, respectively. Interestingly, immature archetype HFD AT 4 was enriched in the middle of the trajectory, followed by archetype *Lep^ob/ob^* AT 3 (**Figure 3G**). Strikingly, the four archetypes within this trajectory displayed increasing *Cck* expression: WT AT 4 was *Cck*-neg, HFD AT 4 and *Lep^ob/ob^* AT 3 were *Cck*-mid, and *Lep^ob/ob^*AT 2 was *Cck*-hi (**Figures 3G** and **3H**), consistent with *Cck* being a key marker of β cell adaptation to obesity.

We next confirmed these findings by *in vivo* lineage tracing analysis. We observed that expression of the transcription factor Neurogenin 3 (Ngn3) was enriched in immature β cells in WT AT 4 and was maintained in the HFD AT 4 and *Lep^ob/ob^* AT 2 populations (**Figure 3C**). Ngn3 drives β cell maturation by binding to the promoter of β cell-specific transcription factors (Pax4 and NeuroD), and its expression is lost once β cells differentiate to a mature phenotype^63^. To label immature β cells, we crossed *Ngn3-Cre^ERT^*^2^ mice^64^ with *Lep^ob/ob^; Rosa26^LSL-TdTomato^* mice and administered tamoxifen at 4 weeks of age before obesity onset (**Figure 4A**). Strikingly, the proportion of insulin+/TdTomato+ comparing 12 weeks vs. 1-week post-tamoxifen increased significantly in obese (*Lep^ob/ob^*) but not lean (*Lep^ob/+^*) mice (**Figures 4E** and **S4E**). These data argue that obesity leads to an expansion of the *Ngn3*+ population and is consistent with the proliferative potential of immature adult β cells. Conversely, the fraction of insulin+/TdTomato+ significantly decreased in obese *Ins1-Cre^ERT^* mice (**Figure 4E**), indicating that mature β cells present prior to obesity onset were replaced. Importantly, neither TdTomato+ nor CCK+ β cells in the *Ngn3-Cre^ERT^*^2^ or *Ins1-Cre^ERT^* lines at 16 weeks of age were dividing (**Figure S6**), arguing that β cell neogenesis had already occurred and that CCK+ cells were largely post-mitotic at this time point. Finally, the majority of CCK+ cells were TdTomato+ in obese *Ngn3-Cre^ERT^*^2^ mice (**Figures 4C** and **4F**) (comparable to *Ins1-Cre^ERT^* despite reduced initial β cell labeling (**Figures 4E** and **S4B**)). These data – concordant with *in silico* analyses (**Figure 3F**) – reveal that an immature subpopulation of postnatal β cells expand and adapt towards a CCK+ state.

### Obesity is associated with β cell transcriptional signatures of endoplasmic reticulum stress

To characterize transcriptional dynamics along the axis of *Cck*-hi *Lep^ob/ob^* AT 2 development, we clustered all highly variable gene trends (see **Methods**) to *Lep^ob/ob^* AT 2 over time into genes that are decreasing (blue) or increasing (red) over the obesity progression (**Figure 5A**). We then performed gene set enrichment analysis (GSEA) for each gene cluster^65,66^ (see **Methods)**. As expected, genes decreasing with obesity were significantly (*q* < 0.05) associated with gene expression regulation in β cells and regulation of insulin secretion (**Figure 5B** and **Table S2A**). Upregulated genes over the obesity trajectory were significantly (*q* < 0.05) associated with diabetes and cellular stress responses including oxidative (electron transport chain, oxidative phosphorylation) and ER stress (**Figure 5B** and **Table S2B**). Several additional gene sets related to the unfolded protein response and protein processing in the ER were also significantly enriched for genes increasing over the progression of obesity (**Table S2B**). Together, these results argue that obesity promotes transcriptional signatures of cell stress and that oxidative and ER stress are prominent factors in the etiology of the obesity-induced stress response.

**Figure 5:**
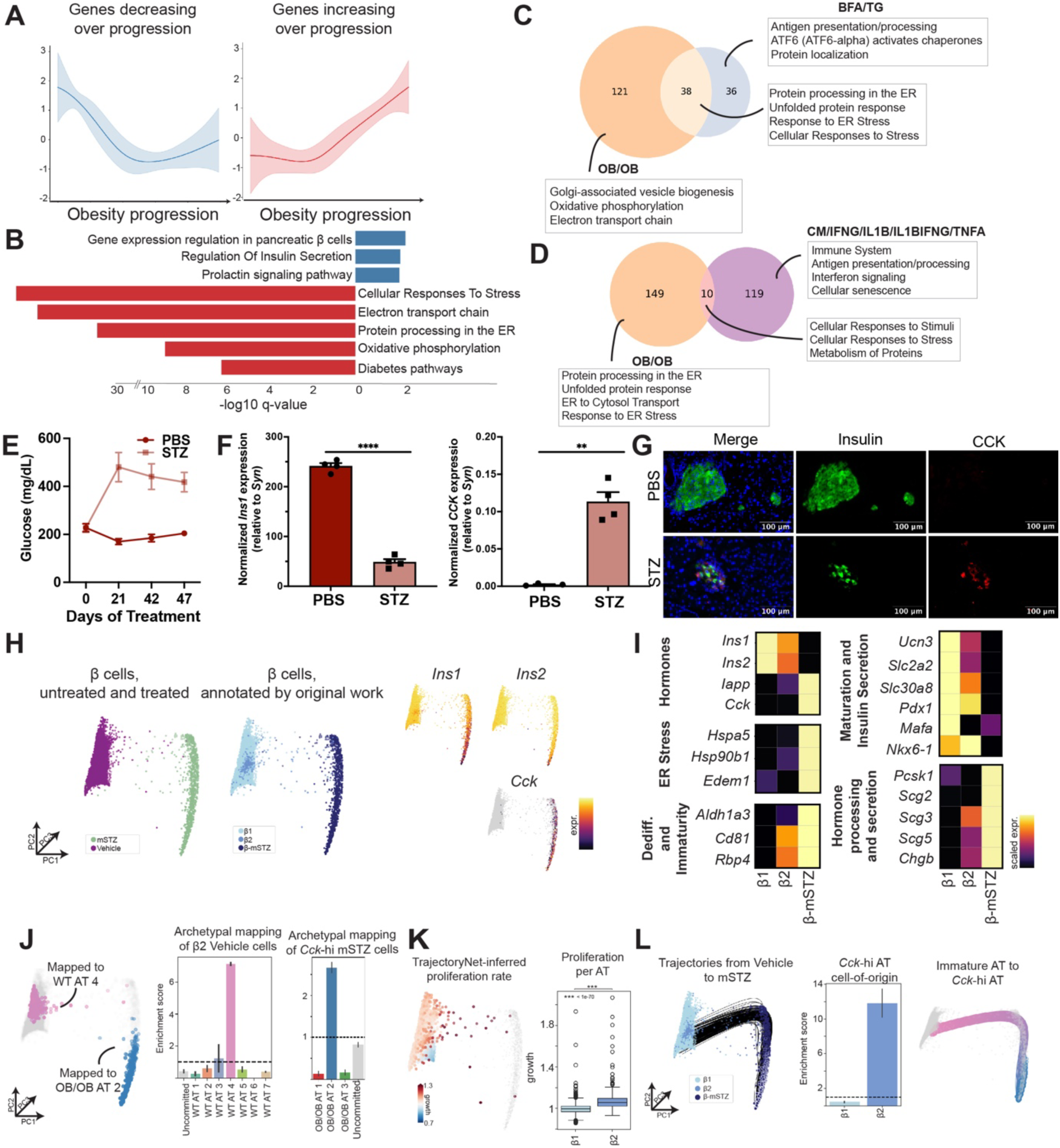
Obesity is associated with transcriptional signatures of increased oxidative and endoplasmic reticulum stress in β cells. (A) Gene clusters derived from gene trends to *Lep^ob/ob^* AT 2, grouped into highly variable genes decreasing and increasing over the obesity progression. (B) Gene set enrichment analysis shows significantly (*q* < 0.05) enriched gene sets for each gene cluster. Red is upregulated. Blue is downregulated. Gene sets and associated genes included: gene expression regulation in β cells (BioPlanet, e.g. *Mafa*, *Slc2a2*, *Neurod1*, *Nkx6.1*), regulation of insulin secretion (GO Biological Process GO:0050796, e.g. *Slc30a8*, *Slc25a22, Cdk16*), cellular responses to stress (Reactome R-HSA-2262752, e.g. *Fos*, *Dnajc3*, *Hspa1a*, *Hsp90b1*), electron transport chain (BioPlanet, e.g. *Ndufb9*, *Slc25a4*, *Atp5*), oxidative phosphorylation (BioPlanet, e.g. *Cox4I1, Atp5*), diabetes pathways (BioPlanet, e.g. *Slc30a5*, *Pdia5/6*, *Dnajc3*), and response to ER stress (GO Biological Process, GO:0034976, e.g. *Pdia3*, *Jun*, *Selenok*, *Hsp90b1*, *Dnajb9*) (C) Gene set enrichment overlap with primary human islet cells exposed to pharmacologic inducers of ER stress (*p* = 1.21×10^-48^, hypergeometric test). (D) Gene set enrichment overlap with primary human islet cells exposed to cytokine inducers of inflammatory stress (*p* = 7.88×10^-5^, hypergeometric test). (E) Serial random glucose (mean +/- s.e.m., n=4 mice/group) of C57/B6 male mice treated with STZ or PBS (vehicle). (F) Final pancreatic *Ins1* and *Cck* expression (normalized to synaptophysin (*Syn*) as general islet cell marker) by qRT-PCR in mice in (**E**). \*\**p*<0.01, **** *p*<0.0001, Welch’s t-test. (G) Immunofluorescence demonstrates loss of insulin and increased CCK protein levels in islets of STZ-treated vs. PBS-treated mice in (**E**). (H) Embedding of β cells from vehicle and mSTZ-treated conditions, colored by treatment, cluster annotation from original work^56^, and marker genes. (I) Heatmap of row-normalized marker gene expression for three β cell subclusters in (**H**). (J) Visualization of vehicle and mSTZ cells mapped to WT AT 4 and *Lep^ob/ob^* AT 2. Enrichment scores (mean +/- 95% confidence interval, n=2 scMMGAN runs) for each WT archetype for β2 vehicle cells and each *Lep^ob/ob^* archetype for *Cck*-hi mSTZ cells. (K) TrajectoryNet-inferred proliferation rate calculated for vehicle cells is significantly higher for β2 than β1. Box plots display 25^th^, 50^th^, and 75^th^ percentiles +/- 1.5 interquartile range (IQR). \*\*\**p* < 1×10^-70^, Wilcoxon rank sum test. (L) Visualization of trajectories from vehicle to β-mSTZ cells mapped to *Lep^ob/ob^*AT 2. The cell-of-origin enrichment score (mean +/- 95% confidence interval, n=2 TrajectoryNet runs) is higher for β2 than β1.

To confirm these findings in orthologous systems, we compared our data to recently published transcriptional signatures derived from primary human islets treated with various β cell stress agents^67^ including drugs that induce ER stress (brefeldin A (BFA) and thapsigargin (TG)) and inflammatory stress-inducing cytokines (IFNɣ, TNFɑ, and/or IL1β). Enriched gene sets in the BFA- and TG-treated (vs. control) islets – including those associated with ER stress – highly and significantly overlapped with gene sets enriched over the obesity progression (*p* = 1.21 x 10^-48^, hypergeometric test) (**Figure 5C** and **Table S3**). While human islets treated with cytokines showed significant overlap with the obesity signatures (*p* = 7.88 x 10^-5^, hypergeometric test), the overlapping gene sets largely represented a general stress response (**Figure 5D** and **Table S3**). These data suggest that obesity induces transcriptional signatures of ER stress in β cells in mice.

We further analyzed the effects of pharmacologic β cell stress induction *in vivo* in mice subject to multiple low doses of streptozotocin (STZ)^56^. STZ is taken up specifically in β cells via the *Glut2* transporter and induces metabolic (NAD+ depletion) and ER stress^68–71^. Like obesity, STZ treatment led to hyperglycemia, decreased pancreatic *Ins1* expression, and increased β cell *Cck* expression in mice (**Figures 5E-G**). To determine the molecular mechanisms of STZ-induced *Cck* expression, we reanalyzed a previously published scRNA-seq dataset of β cells from mice treated with STZ vs. vehicle control^56^. Visualization of these data showed distinct β cell clusters corresponding to treatment groups (β-mSTZ for STZ-treated mice), and two refined subclusters (β1 and β2) within the vehicle control β cell cluster, as previously described^56^. Concordant with obesity, *Ins1/Ins2* expression decreased and *Cck* expression increased with STZ treatment (**Figure 5H**). β1 cells displayed high expression of maturation markers and β cell identity transcription factors (TFs) (**Figure 5I**). In contrast, β2 cells reflected an immature state in vehicle-treated mice, with decreased expression of insulin secretion and maturation genes. β-mSTZ cells were reminiscent of *Lep^ob/ob^* AT 2 cells, exhibiting loss of β cell maturation markers but increased *Cck* and dedifferentiation/immaturity, ER stress, hormone processing, and secretion markers (**Figure 5I**).

Given this similarity, we sought to map β1, β2, and β-mSTZ cells directly to β cells in our obesity dataset for comparison across all transcriptional measurements. However, due to systematic technical variation from differences in handling cells in distinct batches (batch effects), the datasets were not directly comparable. To overcome this, we employed scMMGAN^26^, a data integration approach that uses adversarial learning to align cells from different batches based on their data geometry. scMMGAN enables batch integration and preserves biological variation for direct comparison across diverse experimental setups and stressors. We used scMMGAN to learn the best mapping of vehicle-treated β cells (β1, β2) to WT β cell archetypes and β-mSTZ cells to *Lep^ob/ob^* β cell archetypes using our trained AAnet models (**Figure 5J** and **Table S4**, see **Methods**). The archetypal mapping of the vehicle-treated β cells revealed representation of all WT archetypes, where the immature β2 state was enriched for immature WT AT 4 (**Figure 5J**). Additionally, β-mSTZ cells also displayed representation of all *Lep^ob/ob^* archetypes, where *Cck*-hi (>95^th^ percentile expression) STZ β cells were enriched for *Lep^ob/ob^* AT 2 (**Figure 5J**). Based on results in our obesity models (**Figure 3F**), we hypothesized that *Cck*-hi β-mSTZ cells originate from immature β2 cells. To test this, we ran TrajectoryNet to trace back the most likely paths from β-mSTZ cells assigned to *Lep^ob/ob^* AT 2 to the vehicle-treated β cells. As expected, TrajectoryNet-learned relative proliferation rates determined that β2 cells have a significantly higher growth rate than β1 cells (**Figure 5K**). TrajectoryNet further demonstrated that β2 cells – rather than β1 – were far more enriched within the cells-of-origin for *Cck*-hi β-mSTZ cells (**Figure 5L**). These data argue that both physiologic (obesity) and pharmacologic (STZ) stressors induce immature β cell adaptation to a *Cck-hi* state *in vivo*.

### The obesity progression in mouse latent-space archetypal analysis correlates with type 2 diabetes in humans and mice

We next evaluated whether obesity mirrored other physiologic perturbations of β cell function, including age and type 2 diabetes (T2D). Specifically, we used scMMGAN to map β cells from previously published non-diabetic (ND) and T2D mouse scRNA-seq datasets^35^ totaling 85,129 cells from 31 datasets integrated across conditions and developmental stages (**Figure S7A** and **Table S4**). Along the obesity progression axis, *Ins1* and *Ins2* expression from the mouse reference cells decreased, and *Cck* expression increased (**Figure S7A**). Predominantly, cells from T2D models (*db/db* and mSTZ) mapped to the latter end of the obesity progression, and cells from normal mice – including aged (4m and 2y) mice – mapped to the WT region of the obesity progression (**Figure S7B**). These data suggest that diabetes induction, and not age, aligns to β cell adaptation in obesity. Though there were few analyzed β cells from prenatal mice (Theiler stages 20, 22, and 23), these cells mapped to β cells further along the obesity progression, arguing that obesity induces β cell dedifferentiation to an embryonic state.

To further refine this β cell mapping using our trained AAnet models, we annotated the archetypal assignment of each cell and calculated the proportion of each archetype for all datasets (**Figure S7C**). All archetypes were represented in at least one ND dataset and at least one T2D dataset, and most archetypes were represented in multiple datasets. Additionally, datasets from the same condition showed highly similar archetypal proportions. T2D datasets were predominantly enriched for *Lep^ob/ob^* archetypes (**Figure S7C**). Embryonic ND samples had a significantly higher proportion of *Lep^ob/ob^* AT 2 versus other samples (**Figure S7D** and **S7E**), suggesting *Cck*-hi cells result from dedifferentiation towards an embryonic state. In contrast, postnatal P16 mice (when β cells begin to mature) had a significantly higher proportion for WT AT 4 (**Figures S7D** and **S7E**), an immature state.

To determine the relevance of our findings to human biology, we used scMMGAN to map five published human scRNA-seq datasets consisting of 15 ND and 9 T2D donors to all cells from WT, HFD-fed, and *Lep^ob/ob^*mice (**Table S4)**. The mapped insulin (*INS*) expression for human donors decreased along the obesity trajectory, recapitulating β cell changes along the mouse obesity progression (**Figure 6A**). We annotated the archetypal assignment of each cell for each donor and calculated the proportion of each archetype (**Figure 6B**). All archetypes were represented in at least one ND sample, and all archetypes (except WT AT 2) were represented in at least one T2D sample. Most archetypes were present in multiple patients. When comparing archetypal proportions between ND and T2D donors, mature β cells (WT AT 1) were notably more abundant in ND donors. In contrast, *Lep^ob/ob^*AT 2 and *Lep^ob/ob^* AT 3 cells were present in significantly greater proportions among T2D donors (**Figure 6C**). To further assess the relationship between measured clinical variables in human donors (age, sex, body-mass index (BMI), and T2D status) and the obesity progression axis in our dataset, we leveraged the patient embedding approach DiffusionEMD^27^. As clinical variables are determined on the level of each donor, rather than each individual cell, it is not sufficient to compare the relationship between each variable and the cellular progression axis. DiffusionEMD overcomes this problem by using optimal transport to map an embedding of donors based on the distribution of each donor’s cells on the obesity progression. That is, donors with similar cell distributions are embedded close together, and donors with very different cell distributions (*e.g.*, opposite sides of the obesity progression) are embedded far apart. The resulting DiffusionEMD embedding revealed a linear trajectory across donors, where donors on the left are mapped early on the obesity progression (high *INS*), and donors on the right are mapped late on the obesity progression (low *INS*) (R=-0.93) (**Figure 6D**). Visualizing clinical variables showed a low positive to negligible correlation between age (R=0.28), BMI (R=0.18), and sex (R=0.17) (**Figure 6E**) with the obesity progression axis. Instead, we observed much stronger correlations between the obesity progression and HbA1c (R=0.50) and T2D status (R=0.49) (**Figure 6E**). These results suggest that the β cell adaptations associated with obesity progression in mice are concordant with T2D across species, donors, and scRNA-seq datasets.

**Figure 6.**
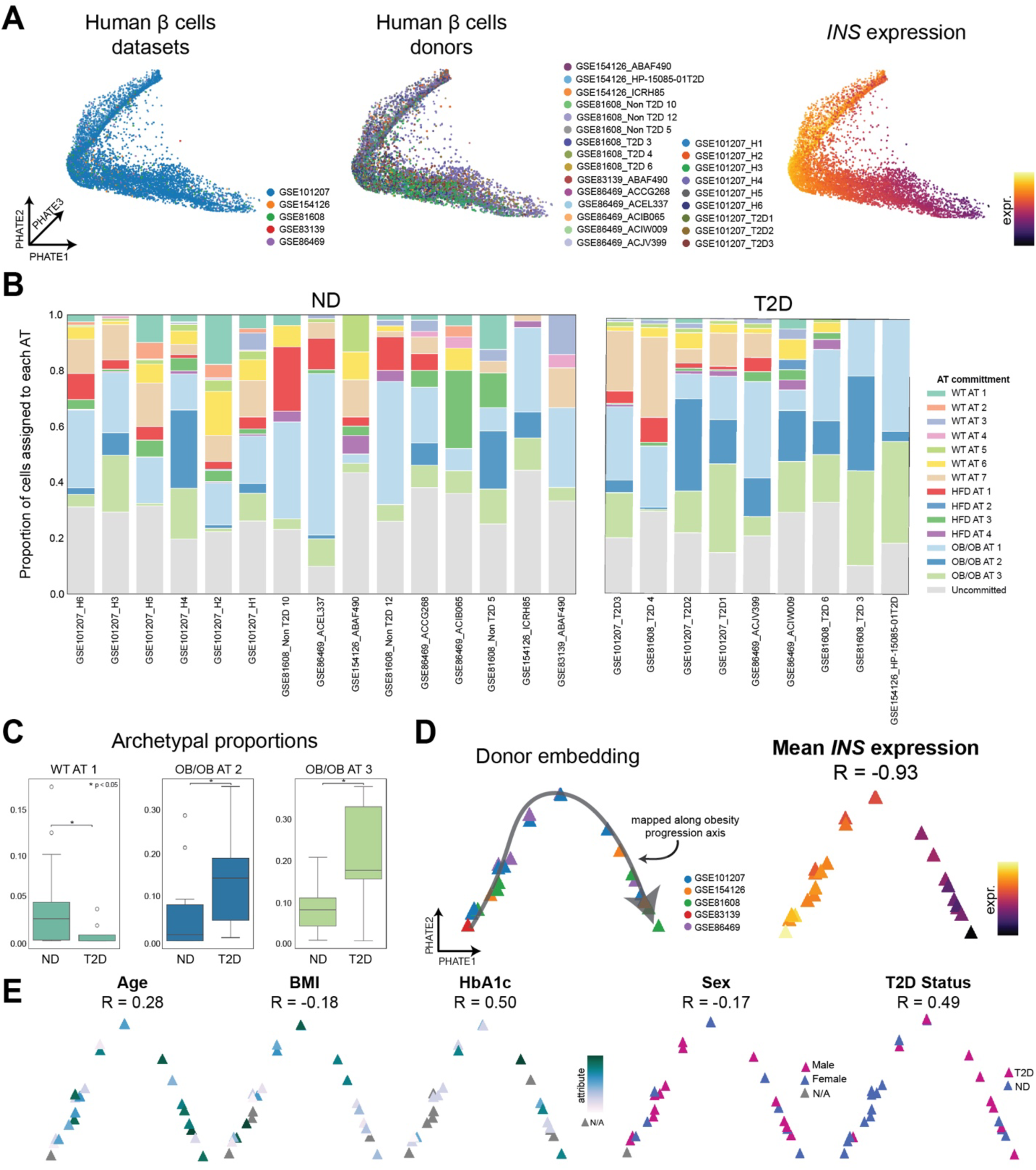
Obesity progression in mouse latent-space archetypal analysis correlates with human T2D. (A) Embedding of human β cells from non-diabetic (ND) and type II diabetes (T2D) donors, mapped onto the obesity progression with scMMGAN. Embedding colored by sample datasets, individual donors, and mapped *INS* expression. (B) Proportion of each archetype for each ND and T2D dataset. (C) Archetypal proportion is significantly different between ND and T2D samples in (**B**) for WT AT 1 (higher in ND samples), *Lep^ob/ob^* AT 2 (higher in T2D samples), and *Lep^ob/ob^* AT 3 (higher in T2D samples). Box plots display 25^th^, 50^th^, and 75^th^ percentiles +/- 1.5 interquartile range (IQR). \**p* <0.05, Wilcoxon rank sum test. (D) Embedding of patient samples, mapped to the obesity progression axis, where each triangle corresponds to an individual donor. Axis direction determined by mean mapped *INS* expression of donors. (E) Fiedler vector of patient sample embedding is colored by age, BMI, HbA1c, sex, and T2D status of donors. Gray corresponds to data not available. Correlation with the murine obesity progression is shown.

### JNK/cJun signaling modulates CCK expression in β cells

Finally, we sought to determine the molecular mechanisms underlying β cell *Cck* expression in obese mice, which might mediate the pro-tumorigenic β cell phenotype. TFs previously associated with *Cck* regulation predominantly decreased in expression along the obesity trajectory except for *Jun*, *Fos*, and *Crem*, which increased (**Figure 7A**). To predict which TFs regulate *Cck* expression, we leveraged our prior approach^23^ which first computes Granger causality scores between highly variable TFs and genes and subsequently prunes to TF-target interactions with high Granger causality and prior evidence for transcriptional regulatory interactions in a reference database^72^ (see **Methods**). We then subsetted to the top 100 regulatory TFs (based on Granger causality scores over all targets) and their targets with high Granger scores (**Table S5**). For visualization, we further subsetted to only TF-target interactions where the TF and the target both increased in expression along the obesity progression trajectory. This resulted in a subnetwork of 39 interactions across 36 TFs and targets, including *Cck* (**Figure 7B**). The subnetwork was enriched (*q* < 0.05) for the AP-1 transcription factor network, including *Jun* and *Fos* directly connected to *Cck*, as well as *Egr1*, *Act1*, *Ccnd1*, and *Cdk1* within the subnetwork (BioPlanet). CCKR signaling map (Panther P06959, *e.g., Cck*, *Jun*, *Fos, Clu*, *Itgb1*, and *Ccnd1)* was also significantly enriched (**Table S6**).

**Figure 7.**
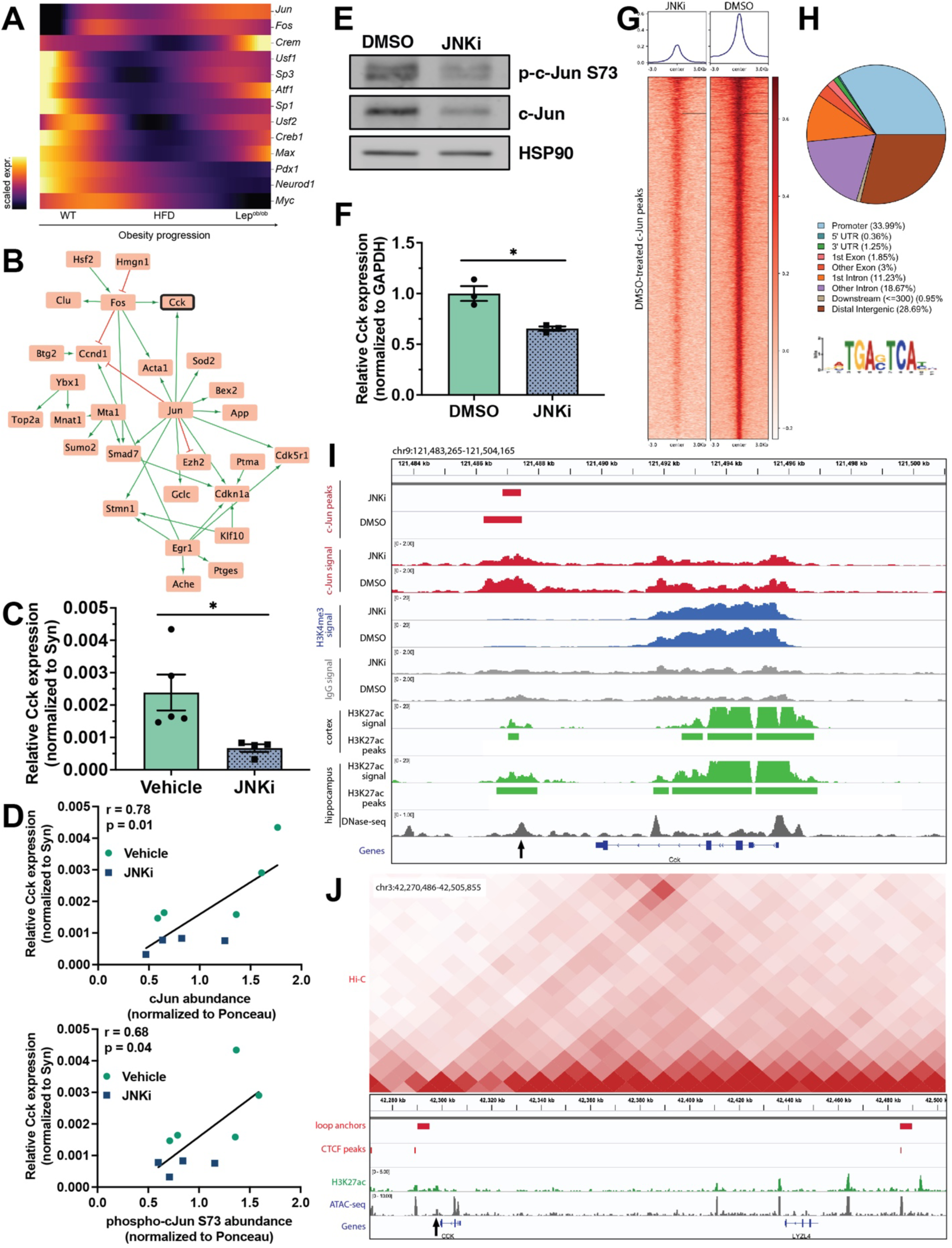
JNK/cJun activation under β cell stress leads to CCK expression. (A) Gene trends to *Cck+ Lep^ob/ob^* cells for putative CCK transcription factors shows a near continuous increase in cJun expression. (B) *In silico* gene regulatory network derived by Granger causality analysis of the TrajectoryNet trajectories. Green edges correspond to activation, and red edges correspond to inhibition. (C) Relative pancreatic *Cck* expression (mean+/-s.e.m., n=4-5 mice/group) by qRT-PCR (normalized to synaptophysin a marker of islet cells) of *Lep^ob/ob^* mice treated with JNKi (20 mg/kg SP-600125 x5 days) or vehicle control. * *p*<0.05, Welch’s test. (D) Correlation of pancreatic *Cck* expression with cJun and phospho-cJun S73 levels (normalized to total protein (Ponceau)) in *Lep^ob/ob^* mice in (**C)**. Pearson correlation coefficients and *p*-values are shown. (E) Representative western blot of total cJun and phospho-cJun S73 in Min6 cells treated with JNK inhibitor (JNKi, 20 μM SP600125) or control (DMSO) for 48 hours. HSP90 is loading control. (F) Relative *Cck* expression (mean+/-s.e.m., n=3 biologic replicates) by qRT-PCR (normalized to *Gapdh*) of Min6 cells in (**E**). * *p*<0.05, Welch’s test. (G) Aggregation heatmap plots of cJun CUT&RUN in Min6 cells treated with JNKi or DMSO control. cJun signal is normalized to E. coli spike-in and IgG. cJun signal is centered on cJun peaks from DMSO-treated cells. (H) Proportion of cJun binding in annotated genomic regions and enrichment at the consensus AP-1 binding motif (27.2%, *p* = 5.4 x 10^-18^) (I) CUT&RUN tracks showing decreased cJun binding downstream of the *Cck* gene with JNKi (arrow). This cJun peak aligns with a putative mouse enhancer region (DNA hypersensitivity and H3K27Ac ChIP-seq peaks). cJun binding at the *Cck* promoter is not significantly altered with JNKi. (J) H3K27ac ChIP-seq and ATAC-seq show a conserved putative enhancer downstream of *CCK* in human islets (arrow). Hi-C demonstrates that the *CCK* putative enhancer occurs at the boundary of a chromatin loop denoted by loop anchors and CTCF bindings sites.

*Jun* encodes the TF cJun, which is activated and stabilized by phosphorylation by cJun N-terminal kinases (JNK). JNK/cJun signaling is a stress-responsive mitogen-activated protein kinase (MAPK) pathway that can be induced by oxidative and ER stress^73,74^, arguing that β cell Cck transcription may be regulated by JNK/cJun signaling *in vivo*. Consistent with this hypothesis, JNK inhibition (JNKi) decreased pancreatic *Cck* expression in *Lep^ob/ob^* mice proportional to the level of total and phosphorylated cJun (**Figures 7C** and **7D**). Similarly, we observed decreased *Cck* expression in Min6 insulinoma cells treated with JNKi *in vitro* (**Figures 7E** and **7F**). To determine whether cJun directly regulates *Cck* expression, we performed cJun CUT&RUN analysis in Min6 cells, which revealed a reduction in overall peak number and intensity with JNKi treatment (**Figures 7G**), consistent with reduced cJun levels (**Figure 7E**). As expected, cJun principally bound promoters or distal intergenic regions, and the most frequent binding motif matched the canonical AP-1 binding site (5’-TCA[GC]TCA-3’) (**Figure 7H**). cJun was enriched at a region ∼3 kb downstream of the 3’ end of the *Cck* gene, the peak signal intensity of which was selectively reduced with JNKi (**Figure 7I**). Data from ENCODE^75,76^ derived from *Cck*-expressing tissues (mouse cortex and hippocampus) suggested that this region may be an enhancer marked by H3K27Ac and DNA hypersensitivity (**Figure 7I**). Furthermore, analyses of human islet ATAC-seq^77^, H3K27Ac ChIP-seq^77^, and Hi-C^78^ data confirmed that this putative enhancer region is conserved in humans and resides at the boundary of a chromatin loop (**Figure 7J**). This higher-order chromatin structure may provide an additional level of transcriptional regulation of CCK. These data indicate that *Cck* expression is mediated by stress-responsive JNK-cJun signaling via a novel conserved 3’ distal enhancer.

## DISCUSSION

Although other cellular components of the pro-tumorigenic PDAC microenvironment – immune, fibroblast, endothelial, and nerve cells – have received greater attention^79^, β cells are emerging as key drivers of PDAC progression. Here, we show that genetic ablation of β cells (*Akita*) slows tumorigenesis even in the rapidly progressive *KPC* PDAC model (**Figure 1**). Our results – based on more faithful genetic PDAC models – mirror findings in Syrian hamsters in which pharmacologic β cell ablation limits carcinogen-induced (N-nitroso-bis (2-oxopropyl) amine (BOP)) PDAC development^80^. They further confirm a conserved basal role of β cells in exocrine tumorigenesis. Our findings are consistent with recent studies demonstrating that reducing insulin (via partial *Ins1/2* knockout) and endocrine-exocrine insulin signaling (via acinar-specific *InsR* knockout) impedes oncogenic *Kras*-driven tumorigenesis^13,14,81^. Moreover, we disentangle the complex interplay between hyperglycemia, β cell function, and cancer. Specifically, we found that hyperglycemia was insufficient to promote tumorigenesis in *KPC-Akita* mice (**Figures 1C** and **1D**). Human PDAC tumors are largely FDG-PET avid, consistent with efficient glucose uptake^82^, and glucose metabolism is thought to be important for PDAC progression^83,84^. Nonetheless, our data argue that excess glucose – in the absence of β cells – is not a primary driver of PDAC development, lending support to the importance of β cells in integrating host metabolic features to drive pancreatic tumorigenesis.

We focused our study on how β cells adapt to pro-tumorigenic host metabolic states induced by diet and obesity, key risk factors for PDAC development. We previously showed that obesity induces a pro-tumorigenic β cell phenotype marked by the expression of CCK, which in turn was sufficient to promote *Kras*-driven tumorigenesis in mice^18^. Here, we found that *Cck* and insulin inversely correlate in expression with increasing obesity (**Figure S3**). CCK is a survival factor for β cells against diverse physiologic and pharmacologic stresses (obesity, STZ, cytokines), including those that increase insulin demand^85–88^. Its upregulation and the concurrent downregulation of insulin may be protective mechanisms analogous to what is observed with acinar cells under stress. Despite distinct physiologic functions, the endocrine and exocrine compartments of the pancreas adopt convergent paths in handling cellular stresses by downregulating their major protein products (insulin for β cells and proteases for acinar cells) and converting cellular states (β cell dedifferentiation and acinar-to-ductal metaplasia). Importantly, both these cellular transitions serve to augment PDAC risk through dysregulated pro-tumorigenic hormone secretion (*e.g.,* CCK) or by generating a permissive state (ADM) for oncogenic *Kras*-driven transformation^16,18,21,89,90^. Thus, self-preservation of the pancreas may pose an unanticipated threat to the organism at large and speaks to the unique interconnected nature of the endocrine and exocrine compartments and their role in cancer pathogenesis.

Despite their inverse expression trajectories, our data suggest that insulin and CCK reside in the same secretory granules (**Figure S3E**), and therefore are likely to be co-secreted and capable of signaling to nearby exocrine cells. We had previously shown that β cell-specific *Cck* overexpression stimulates acinar cell proliferation *in vivo*^18^. Others have described that peri-islet acinar cells have different gene expression signatures than those found further from the islet in obese *Lepr^db/db^*mice that also exhibit β cell *Cck* expression^18,40^. Importantly, 9 of 15 CCK-regulated genes (vs. 3 of 11 insulin-regulated genes) were upregulated in peri-islet acinar cells assessed by smFISH^40^, including several genes (proteases, *Reg* genes) previously implicated in PDAC pathogenesis^45,91^. Our single-cell analyses confirmed that peri-islet acinar cells adopt altered transcriptional cell states in obesity, that these states may be governed by CCK signaling, and that they are associated with increased islet proximal exocrine tumorigenesis (**Figure 2**). These data argue that CCK may play a more consequential or complementary role to insulin in local acinar cell transformation in the context of obesity.

Our study also provides new insights into the cellular and molecular mechanisms that govern β cell adaptation to obesity and lead to a pro-tumorigenic CCK+ state. We leveraged a novel suite of computational methods (TrajectoryNet, AAnet, scMMGAN, DiffusionEMD) for scRNA-seq data analysis to characterize β cell heterogeneity and transcriptional dynamics with increasing obesity (**Figure S1**), which can promote pancreatic tumorigenesis. These approaches, used in conjunction, represent a significant advance in single-cell analysis to attain novel biological insights, especially in the context of β cell biology. Standard practices in single-cell analysis characterize cellular states based on clustering and pseudotime inference^38^. Indeed, the majority of published single-cell analyses of mouse and human pancreatic islets in normal physiology and diabetes involved unsupervised clustering to identify endocrine and exocrine cell types, then subclustering or identifying factors of variation within cell types of interest, including β cells^92–96^. However, these analyses are limited in their ability to infer cells-of-origin, gene regulatory relationships, and transcriptional dynamics within and across cell types, analyses enabled by TrajectoryNet through calculation of cell-specific trajectories. Furthermore, unlike subclustering, latent-space archetypal analysis (AAnet) avoids arbitrarily discretizing the continuous state space and losing gene signals and behavior that result from high plasticity and transformation between states^25^. In addition, by mapping published datasets directly to our obesity progression through batch integration, scMMGAN established that β cells in the context of obesity are highly related to diverse stressors across the entire transcriptome, beyond marker genes or β cell-specific signatures. Finally, while prior work has shown the relationship between marker gene expression and measured clinical variables^92^, DiffusionEMD instead characterized broad similarity between patients based on all β cell transcriptomic measurements, enabling more accurate correlations of the murine obesity progression with clinical variables. Our single-cell analytic pipeline therefore enables avenues of β cell characterization not possible with existing methodologies.

Using complementary *in silico* and *in vivo* experimental approaches (**Figures 3** and **4**), we found that obesity promotes β cell neogenesis via β cell duplication rather than transdifferentiation of other cell types (*e.g.*, α or duct cells)^51,52^. This finding aligns with the preponderance of evidence that β cell regeneration and islet expansion in physiologic (homeostasis, pregnancy, now obesity) and injury states largely occur through β cell replication^48^ with the exception of non-physiologic complete β cell ablation^49,50^. Leveraging the unique capabilities of AAnet to discover rare subpopulations within heterogenous β cells, we further showed that *Cck*+ β cells largely arise from a single archetype (WT AT 4). This archetype represents an immature β cell population (maps to P16 β cells, lacks maturity TFs) that resembles a recently described regenerative *Ucn3*-/*Ngn3*+/*Ins1/2-*lo immature β cell type that persists into mouse adulthood^97,98^. By lineage tracing *Ngn3*+ cells prior to and following obesity induction, we found that obesity is associated with expansion of this population into β cells that express CCK (**Figure 4**). Our findings in a genetic model of obesity are consistent with a high-fat fast mimicking diet that induced expansion of *Ngn3*+ cells to comprise ∼50% (vs. ∼20% of *ad libitum* feeding; nearly identical to our findings in *Lep^ob/ob^* mice (**Figure 4E**)) of the islet population at 14 weeks of age^61^, arguing that this effect occurs independent of model or diet.

Given CCK’s pro-survival function on β cells^85–88^, it is not surprising that our results suggest that *Cck* induction occurs as a response to both physiologic (obesity) and pharmacologic (STZ) stress. Transcriptional profiling supports specific stressors in *Cck* induction, as the obesity-induced β cell state overlaps more strongly with human β cells under ER rather than cytokine stress (**Figure 5**). These data are consistent with prior observations showing downregulated cytokine production in islet macrophages in the context of both diet-induced and genetic obesity in mice^18,99^. Instead, obesity and associated insulin resistance lead to hyperglycemia and increased insulin demand, resulting in enhanced ER and oxidative stress^100^. Through these stressors, obesity reverts β cells to a dedifferentiated embryonic-like state that aligns with human and mouse T2D datasets (**Figure 6**), consistent with the known association between T2D and β cell dedifferentiation^101,102^. We further identified stress-responsive MAPK signaling via JNK/cJun in the regulation of *Cck* expression in β cell models (**Figure 7**). Previous work has implicated JNK3/cJun signaling in β cell proliferation in normal postnatal development^103^, suggesting that this signaling pathway may also be co-opted to respond to obesity-associated stress. Finally, we discovered a novel JNKi-sensitive cJun binding peak in a conserved putative 3’ enhancer downstream of the *Cck* gene by which cJun may regulate *Cck* expression in a stress-responsive fashion. Taken together, our work lays the groundwork for understanding the stress and signaling pathways that govern the pro-tumorigenic β cell phenotype, enabling future studies targeting these endocrine β cell pathways as a novel approach for the prevention or interception of PDAC.

## Acknowledgments

We thank the Muzumdar and Krishnaswamy lab members for helpful discussions and feedback; Drs. F. Zhang and S. Chen for technical assistance with AAV-Leptin production; the Yale Center for Genome Analysis (YCGA) for library preparation and sequencing; J. Nikolaus and the West Campus Imaging Core for assistance with confocal microscopy; the Center for Cellular and Molecular Imaging (CCMI) Electron Microscopy Facility for electron microscopy; X. Zhao for assistance with mouse islet isolations, islet dispersion, and intact islet perifusion studies at the Chemical Metabolism Core at Yale University; T. Nottoli and the Yale Genome Editing Center (YGEC) for cryorecovery of mouse embryos; and Drs. T. Jacks, K. Kaestner, A. Lowy, A. Leiter, L. Philipson, H. Zeng, and the Allen Institute for Brain Science for mice. C.G. is a recipient of a National Cancer Institute (NCI) Ruth L. Kirchstein National Research Award (NRSA) for predoctoral students (F31-CA268845) and was funded by the Training Program in Genetics (TPG) training grant (5T32-GM148332). A.V. was funded by a Gruber Science Fellowship. D.C.M. and A.S. are supported by the Yale Medical Scientist Training Program (5T32-GM007205). S.S.A. was supported by a postdoctoral fellowship through the Yale Cancer Biology Training Program (T32-CA193200). J.B.J. is supported by a Conquer Cancer Foundation-American Society of Clinical Oncology Young Investigator Award (YIA) and a Yale Center for Clinical Investigation (YCCI) Scholars Award. C.F.R. was supported by a postdoctoral fellowship through the Yale Cancer Biology Training Program (T32-CA193200) and is supported by an NCI Research Supplement to Promote Diversity in Health-Related Research (R01-CA276108-02S1). R.G.K. is supported by the NIDDK (1R01-DK127637). S.K. is supported by the National Science Foundation (NSF Career Grant 2047856, NSF DMS grant 2327211, and NSF CISE grant 2403317). M.D.M. acknowledges support from an NIH Director’s New Innovator Award (DP2-CA248136), Lustgarten Foundation Therapeutics Focused Research Program (TFRP), NCI (R01-CA276108), and in part, the Yale Comprehensive Cancer Center Support Grant (P30-CA016359). The content is solely the responsibility of the authors and does not necessarily represent the official views of the National Institutes of Health. This work was largely supported by Damon Runyon-Rachleff Innovation Awards (66-21/66S-21) to M.D.M.

## Author contributions

C.G. and M.D.M. conceived of and designed the study. C.G., D.C.M., and S.S.A., A.S., J.B.J., and C.F.R. performed mouse and cell line experiments and associated analyses. A.V. and A.T performed computational experiments and analysis, which were supervised by S.K. R.C. performed islet isolation for scRNA-seq and perifusion experiments, which were supervised by R.G.K. R.S. performed analysis of CUT&RUN data. S.K. and M.D.M. supervised the overall study. C.G., A.V, and M.D.M wrote the manuscript with input from all authors.

## Declaration of interests

M.D.M. and S.S.A. are inventors on a patent applied for by Yale University that is unrelated to this work. M.D.M. received research funding from a Genentech supported AACR grant and an honorarium from Nested Therapeutics. The remaining authors declare no competing interests.

## METHODS

### Animal studies

Animal studies were performed at Yale University West Campus and approved under the Yale University Institutional Animal Care and Use Committee (IACUC) protocol #20170. *Akita* (Stock #003548), *Kras^LSL-G12D^* (Stock #008179), *Lep^ob/ob^* (Stock #000632), *Ins1-Cre^ERT^*(Stock #024709), *Gcg-Cre^ERT^*^2^ (Stock #042277), *Ngn3-Cre^ERT^*^2^ (Stock #028365), *Pdx1-Cre* (Stock #014647), *Rosa26^LSL-TdTomato^* (Stock #007914), and *Trp53^LSL-R^*^172^*^H^* (Stock #008652) mice were obtained from the Jackson Laboratory (JAX). Cryorecovery of *Gcg-Cre^ERT^*^2^ and *Ngn3-Cre^ERT^*^2^ embryos was performed by the Yale Genome Editing Center. *KPC-Akita*, *KCO*, and *Cre^ER^; Lep^ob/ob^; Rosa26^LSL-TdTomato^* mice were generated by breeding. Genotyping of the mice was done via PCR using genomic DNA from the tail or ear as a template. Tail DNA was isolated using Hotshot extraction, and PCR was run using GO Taq Green Mastermix (Promega). Primers and protocols used were obtained from JAX. For *Lep^ob/ob^* breedings, *Lep^ob/ob^* mice were administered AAV2/1-CAG-mLeptin via intramuscular injection once, which restores fertility^18^. For bulk RNA-seq experiments, *Lep^ob/ob^* and congenic age-matched C57/B6 WT mice (Stock #000664) were purchased from JAX. *Lep^ob/ob^*mice were treated with CCK receptor antagonists (proglumide (0.1 mg/mL; Sigma) or lorglumide sodium salt (0.05 mg/mL; Sigma)) or control drinking water continuously for 6 weeks starting at 6 weeks of age. Mice were euthanized at 12 weeks of age and isolated pancreata were snap frozen for RNA-seq analysis, as described below. *Cre^ER^* mice were induced at ∼4 weeks of age with 8mg/40g (obese *Lep^ob/ob^*) or 6 mg/40g (non-obese *Lep^ob/+^*) of tamoxifen (Sigma) in corn oil via oral gavage for 4 consecutive days. Mice were euthanized 1- or 12-weeks following start of tamoxifen administration for analysis. For STZ experiments, male mice were treated with 50 mg/kg per day of STZ or PBS (vehicle) by intra-peritoneal injection for 5 consecutive days and analyzed 7 weeks afterwards. For JNKi experiments, 17-week-old *Lep^ob/ob^* mice were treated with either 20mg/kg SP-600125 (MedChemExpress) or vehicle control (5% DMSO, 40% PEG-300, 5% Tween 80, 50% ddH_2_O) by oral gavage once daily for five days prior to dissection. Mouse weights were measured using a small animal scale, and non-fasting random blood glucose was measured using a OneTouch Ultra2 glucometer. Mice of both sexes were used for all experiments (and denoted in the figures with symbols) except for the STZ and scRNA-seq experiments for which only male mice were used due to reduced β cell effects of STZ and obesogenic effects of HFD, respectively, in female mice.

### Cell culture

Min6 cells (at a passage of 31-50) were grown under high glucose DMEM (Corning) supplemented with 15% fetal bovine serum (FBS; ThermoFisher Scientific) Penicillin streptomycin (1%; ThermoFisher Scientific), L Glutamine (2mM; ThermoFisher Scientific), Sodium Pyruvate (1mM; ThermoFisher Scientific), and 2-mercaptoethanol (0.05mM; Sigma). Cells tested negative for mycoplasma by PCR performed by the Yale Molecular and Serological Diagnostics Core.

### Immunoelectron microscopy

Cultured Min6 cells were fixed in 0.1% glutaraldehyde, 4% paraformaldehyde in 0.1M sodium cacodylate buffer (pH 7.4) for 30 minutes at room temperature, followed by 4% paraformaldehyde in the same buffer for one hour. After buffer rinse, cells were scraped off the culture dish and gently centrifuged. Pelleted cells were resuspended in 10% gelatin and submerged in 2.3M sucrose in PBS at 4°C overnight prior to freezing in liquid nitrogen. 60-nm thick sections were cut using a Leica EM UC6/FC6 cryo-ultramicrotome and collected on nickel grids. After warming to room temperature, sections were immunolabeled with primary antibodies for 30 minutes, followed by 30-minute incubation with immunoglobulin G antibodies coupled to 5 nm and 10 nm protein A-gold particles (Utrecht University Medical Center, Netherlands). Images were acquired using a FEI Tecnai Biotwin transmission electron microscope (80 kV) and an AMT NanoSprint15 cMOS camera.

### Western Blot

Mouse pancreata were snap-frozen in liquid nitrogen and manually pulverized using the BioPulverizer (BioSpec Products 59012MS). Pancreatic tissue (∼40 mg) and Min6 whole cell lysates were harvested in RIPA buffer (Pierce), EDTA (1:100) (ThermoFisher Scientific), and Halt protease and phosphatase inhibitor cocktails (1:100) (ThermoFisher Scientific). Tissue samples were vortexed and homogenized using a pellet mixer (VWR) and rotated at 4°C for one hour prior to centrifugation at maximum speed for 20 minutes. Cells were scraped and placed on a rotator at 4°C for 15 min and then spun as above. Protein concentrations were measured using the BCA Protein Assay Kit (Pierce) following the manufacturer’s protocol. Equal amounts of protein (30 µg) were loaded to each well of a mini-PROTEAN 4-20% TGX stain-free precast gels (Bio-Rad). Protein was transferred to nitrocellulose membranes using the TransBlot Turbo Transfer System (Bio-Rad). For pancreatic tissue lysates, membranes were stained with Ponceau S (0.5% (w/v) in 1% glacial acetic acid) to verify equal protein loading and then washed with PBS-T (PBS, Tween-20 (0.1%)). Membranes were blocked using Odyssey Blocking Buffer (Licor) in PBS (1:2) for one hour, then incubated with primary antibodies diluted in Odyssey Blocking Buffer and PBS-T (1:2) overnight at 4°C. Membranes were washed for 10 minutes with PBS-T 3 times and then incubated in DyLight secondary antibodies (Cell Signaling Technologies) for one hour at room temperature. Membranes were washed with PBS-T for 10 minutes, 3 times each, prior to imaging. Membranes were imaged using the ChemiDoc Touch Imaging System (Bio-Rad). Band intensity was quantified using ImageLab software.

### RNA Extraction, qRT-PCR, and bulk RNA-sequencing

Mouse pancreata were snap frozen in liquid nitrogen and manually pulverized using the BioPulverizer (BioSpec Products 59012MS). RNA was extracted from pulverized tissue (20-40 mg) and Min6 cells, homogenized via QIAshredder column (Qiagen), and purified using the Qiagen Mini RNeasy kit following the manufacturer’s protocol with DNAse I treatment to remove any genomic DNA contamination. RNA quality and quantity was assessed using an Agilent Bioanalyzer, and only high-quality RNA samples (RIN>7) were used in downstream analysis. 1 μg of RNA was used to synthesize cDNA using Applied Biosystems™ High-Capacity cDNA Reverse Transcription Kit. cDNA was diluted in nuclease-free water (1:5), and qPCR was run using TaqMan™ Universal PCR Master Mix and FAM probes (ThermoFisher Scientific) for GAPDH (Mm99999915_g1), Insulin 1 (Mm01950294_s1), Synaptophysin (Mm00436850_m1) and CCK (Mm00446170_m1). qPCR assays were run on a CFX Opus 384 Real-Time PCR System (Bio-Rad). RNA-seq libraries were prepared with polyA selection (Illumina) and sequenced (>25M 100-bp paired-end (2×100) reads on NovaSeq (Illumina)) with the Yale Center for Genome Analysis (YCGA). Low-quality bases and adapter sequences were trimmed using Trim Galore (v0.6.7) and CutAdapt (v3.5). Cleaned reads were aligned to the UCSC mouse genome (mm10) using the STAR aligner (v2.7.9). Differential gene expression analysis was performed using DESeq2 in R, and genes with an adjusted *p*-value < 0.05 were considered differentially expressed.

### CUT&RUN

Min6 cells were plated (2.5 million) and grown for a total of 3 days. The day after plating, cells were treated with the JNK inhibitor SP600125 (20 µM, Med Chem Express) or DMSO (control) for 48 hours. 550,000 cells were harvested per reaction and condition. Cells were processed using the CUTANA ChIC/CUT&RUN Kit (14-1048), JUN/cJun CUTANA CUT&RUN antibody (SKU:13-2019), and positive (anti-H3K4Me3) and negative (IgG) control antibodies from the kit following the manufacturer’s protocol. Library preparation was performed by the YCGA followed by sequencing to a depth of 30 million paired-end 150 bp reads (2×150) per sample on NovaSeq (Illumina).

### Mouse tissue preparation for histology and immunohistochemistry

Mice were euthanized by CO_2_ asphyxiation. Pancreas was isolated, fixed in 10% neutral-buffered formalin overnight, dehydrated in 70% ethanol, and embedded in paraffin by Yale Tissue Pathology Services. Adjacent 5-μm sections were cut and stained with hematoxylin and eosin or subject to immunohistochemistry (IHC) using a ThermoFisher Scientific Autostainer 360. Mach 2 HRP-labeled micro-polymers (Biocare Medical) were used for primary antibody detection. Tissues were imaged with a modified Nikon T2R inverted microscope (MVI), 4x/10x/20x/40x objectives, and a 2.8 MP CoolSNAP Dyno CCD camera (Photometrics). Monochromatic red, green, and blue images were merged using ImageJ software (NIH).

### Mouse tissue preparation for immunofluorescence

Mice were euthanized by CO_2_ asphyxiation and fixed with 4% paraformaldehyde (PFA; Electron Microscopy Sciences) via cardiac perfusion. The pancreas, small intestines, and liver were harvested and fixed in 4% PFA overnight at 4°C, washed in PBS three times for 5 min, and cryoprotected in 30% sucrose in PBS at 4°C. Tissue was embedded in O.C.T. (Tissue-TEK) and stored at -80°C prior to cryosectioning. 10-μm cryosections were obtained using a Leica Cryostat (CM1860). Slides were stored at -80°C until immunofluorescence staining. For immunofluorescence, slides were thawed at room temperature for one hour and washed three times for 10 min with PBS. Tissue was permeabilized and blocked for one hour with PBT (PBS, TritonX-100 (0.3%)) with 10% donkey serum (Jackson ImmunoResearch). Slides were incubated at 4°C with primary (Day 1) and secondary (Day 2) antibodies (Jackson ImmunoResearch) overnight in PBT and 5% donkey serum in a humidified chamber and washed three times for 10 min after each antibody incubation with PBT. On Day 3, slides were stained with DAPI (1:500; ThermoFisher Scientific) in PBS and washed once with PBS. Slides were mounted using Vectashield Hardset Antifade Mounting Medium (VectorLabs). Slides were imaged at the Yale West Campus Imaging Core using a Leica SP8 Spinning Disk Confocal Microscope and an Andor Benchtop Confocal Microscope BC43.

### Immunocytochemistry

Min6 cells were grown in a 6-well plate with a 20 mm round cover glass (Cell Treat, Part#: 229173) for 3 days, washed with PBS (2x, 5 min each), fixed with 4% PFA (15 min), and washed with PBS (2x, 5 min each) at room temperature. Cells were blocked using PBT with 10% donkey serum and incubated overnight at 4°C in primary antibody in PBT with 2.5% donkey serum. On Day 2, cells were washed with PBS (3x, 5 min each) and incubated overnight at 4°C in secondary antibody in PBT with 5% donkey serum. On Day 3, cells were washed with PBT (3x, 5 min each), incubated with DAPI (1:500) for 5 min, washed in PBS for 5 min, and mounted on slides using Vectashield Hardset Antifade Mounting Medium. Slides were scanned using Yale West Campus Imaging Core’s Leica SP8 Spinning Disk Confocal Microscope.

### Antibodies

| **Antibody Target** | **Species** | **Company** | **Product Number** | **Application** | **Dilution** |
| --- | --- | --- | --- | --- | --- |
| Cholecystokinin | Rabbit | Immunostar | 20078 | IF/ICC/EM | 1:100/<br>1:100/<br>1:250 |
| Insulin | Rat | R&D Systems | MAB1417 | IF/ICC/EM | 1:500/<br>1:500/<br>1:250 |
| Glucagon | Rabbit | Sigma Aldrich | SAB4501137 | IF | 1:400 |
| Ki67 | Rabbit | Cell Signaling Technology | 9129S | IF | 1:100 |
| Phospho-c-Jun (Ser73) | Rabbit | Cell Signaling Technology | 3270S | WB | 1:500 |
| cJun | Rabbit | Cell Signaling Technology | 9165S | WB | 1:1000 |
| Hsp90 | Rabbit | Cell Signaling Technology | 4877S | WB | 1:5000 |
| JUN/cJun | Rabbit | EpiCypher | 13-2019 | CUT&RUN | 5 µg |
| Insulin | Guinea Pig | Accurate Chemical | BMAT5014 | IHC | 1:300 |
| BiP | Rabbit | Cell Signaling Technology | 3177 | IHC | 1:400 |
| Synaptophysin | Rabbit | ThermoFisher Scientific | RB-1461 | IHC | 1:100 |
| Anti-Rabbit IgG (DyLight 800 4X PEG conjugate) | Goat | Cell Signaling Technology | 5151S | WB | 1:2500 |
| Anti-Rat IgG Alexa Fluor Plus 488 | Donkey | ThermoFisher Scientific | A48269 | IF/ICC | 1:500 |
| Anti-Rabbit IgG Alexa Fluor Plus 647 | Donkey | ThermoFisher Scientific | A32795 | IF/ICC | 1:500 |
| Mach2 Rabbit HRP-Polymer | Goat | Biocare Medical | RHRP520 | IHC | 1:1 |
| Peroxidase AffiniPure Anti-Guinea Pig | Donkey | Jackson ImmunoResearch | 706-035-148 | IHC | 1:500 |

### Pancreatic islet isolation, perifusion, and single-cell RNA sequencing (scRNA-seq)

Male 16-week-old wild-type (WT) C57/B6 mice (Stock #000664) and male C57/B6 mice fed a 60% kcal fat diet (Research Diets 12492) for 10 weeks (Stock #380050) starting at 6 weeks of age were obtained from JAX. Islets were isolated, pooled for each group (n=4 mice per group), and processed for perifusion for GSIS measurements or dispersed into single cells for sequencing, as previously described^18^. For scRNA-seq, dead cells were removed using a Dead Cell Removal Kit (Miltenyi Biotec, 130-090-101). Cell concentration and viability were determined using a Countess II Automated Cell Counter (ThermoFisher Scientific). 10X Genomics Chromium Single Cell 3’ Reagent Kit v3 library preparation and sequencing on Illumina HiSeq were performed by the YCGA.

### Quantification and statistical analysis

#### Tumor proximity to islets

Scanned H&E sections were digitally analyzed using QuPath v0.4.3 from lean 13-week-old *KC; Lep^ob/+^* and obese 7-week-old *KC; Lep^ob/ob^* mice (n=4-5 mice/group) in blinded fashion. Tumors were largely low-grade PanINs prior to further tumor progression in these models at the designated timepoints. All low-grade PanINs in the splenic lobe of the pancreas (where most islets are located in mice) of each mouse were quantified and classified as islet proximal or distant. Islets (diameter >100 µm) were defined as being proximal if they resided within a distance of 400 µm from tumor epithelial cell edge to the center of an islet (where β cells are located). This represents the approximate length of ten acinar clusters within a lobule, which is twice the expected diffusion limit but would allow hormone delivery via the interconnected islet-acinar capillary network^104,105^. Fisher’s exact test was performed to compare lean and obese conditions with a significance threshold of *p*<0.05. Despite harboring fewer overall tumors (*n*=58 vs. 146) at this earlier timepoint, a greater proportion of tumors in *KC; Lep^ob/ob^* were close to islets.

#### *In vivo* lineage tracing analysis

To analyze sufficient islets at single-cell resolution for each mouse, we obtained widefield fluorescence images using a Nikon Spinning Disk Confocal (CSU-W1) under 40x or 60x oil immersion with laser wavelengths of 405-nm, 488-nm, 561-nm, and 647-nm. Fluorescence channels were merged using ImageJ, and positive cells were counted manually. Briefly, 15-20 islets were analyzed for each 16-week-old experimental mouse per line, scored for presence or absence of TdTomato, insulin, or CCK expression, and each counted cell was marked in ImageJ to avoid double counting. The percentage of CCK+ or insulin+ cells also expressing TdTomato were calculated for each mouse for statistical comparisons between mice. For mouse line validation (5-week analysis timepoint), 10 islets were analyzed per mouse. For *Ins1-Cre^ERT^*and *Ngn3-Cre^ERT^*^2^ mouse lines, the proportion of insulin+ cells harboring TdTomato fluorescence were quantified. For the *Gcg-Cre^ERT^*^2^ line, we calculated the percentage of Gcg+/TdTomato+ cells and divided the result by the total Gcg+ cell population. Due to unequal variance, the Welch’s t-test was used for statistical analysis when comparing groups with a significance threshold of *p*<0.05.

#### Preprocessing of scRNA-seq data

##### Murine obesity models

scRNA-seq data from 12 week-old C57/B6 *Lep^ob/ob^* and WT mice were previously published by our group^18^, and raw data deposited in GEO (GSE137236) were reanalyzed in comparison to new HFD and WT data described above. We combined batches from all four scRNA-seq datasets and removed rare genes (expressed in fewer than 15 cells). We then filtered and L1 normalized cells by library size (maximum library size=15000) and square-root transformed the data. We finally filtered cells with high mitochondrial expression (maximum expression=12.5). All processed cells were visualized with PHATE^106^, with *Ins1*, *Ins2*, *Sst*, *Ppy*, and *Gcg* removed from PHATE calculation due to very high expression affecting cell-cell distances. All cells were then clustered with *k*-means clustering (n_clusters=20), and clusters were annotated based on published mouse islet marker genes^35^. β cells were then subsetted and further filtered to remove outlier populations with large differences in the number of genes expressed per cell, indicating low quality of the cell populations. Endocrine cell clusters were retained for TrajectoryNet, re-embedded, and clustered with *k*-means clustering (*n*_clusters=10, grouped for annotation). For all other downstream analysis, β (*Ins1/Ins2*+ cells totaling 17,336) or acinar (*Cpa1/Prss2*+ cells totaling 252) were retained. For gene cluster analysis, GSEA of gene clusters, and Granger causality analysis, we used highly variable genes defined using default parameters in Scanpy^107^. Endocrine and acinar cell embeddings were generated with PHATE. β cell embeddings were generated by first denoising with MAGIC^108^, then reducing dimensionality with PCA. This enabled visualization of heterogeneity similar to PHATE while maintaining invertibility to the gene space for trajectory and latent-space archetypal analysis.

##### Published datasets

Upregulated (log_2_FC expression>1.5, FDR<0.05) acinar-specific genes from RNA-seq analysis of islets isolated from *Lepr^db/db^* versus WT mice^39^ were extracted from “Table S3” of Egozi et al.^40^. Differentially expressed genes for patients exposed to pharmacologic β cell stressors were extracted from GSE237448^67^. We subsetted to DEGs per stressor that were enriched versus control (CTRL) in the majority of patients (at least 3 of 5). We then computed the gene sets for DEGs per stressor. For BFA and TG, we took the intersection of gene sets before comparing overlap with obesity. For CM, IFN, IL1B, IL1BIFNG, and TNFA, we took the intersection of gene sets before comparing overlap with obesity. All gene sets enriched per stressor and associated genes are provided in **Table S3**.

Pre-processed and annotated scRNA-seq data from islets derived from mice treated with STZ (mSTZ) or control (vehicle) were obtained from GSE211799^56^ and subsetted to β cells. β1, β2, and β-mSTZ annotations were used from original work, except for six control cells labeled as β-mSTZ, which were relabeled as β2 for cell-of-origin analysis to differentiate between an immature pre-existing population and a differentiated cell state.

Human islet datasets (GSE101207, GSE81608, GSE86469, GSE154126, and GSE83139) were harmonized across measured clinical variables, subsetted to β cells from ND and T2D donors, and pre-processed for library size normalization and square-root transformation. The mouse atlas^35^ from was obtained from https://cellxgene.cziscience.com/collections/296237e2-393d-4e31-b590-b03f74ac5070 and subsetted to only β cells from control/ND and type 2 diabetic (T2D) mouse models. Samples with fewer than 2 cells were excluded. Samples annotated “Embryonic” show embryo progression from E12.5 to E15.5. Samples annotated as P16, 4m, and aged were islet cells isolated from 16 day-old, 4-month-old, and 2-year-old mice and sorted according to the Fltp lineage-tracing model. mSTZ samples are derived from healthy adult control mice, an STZ-treated model, and mSTZ models subject to different anti-T2D treatments. *db/db* samples are from healthy adult control, *Lepr^db/db^*, and *Lepr^db/db^* subject to different anti-T2D treatments. Chem samples are healthy young adult control samples (only control retained for this analysis).

#### TrajectoryNet

##### Overview

Traditionally, characterizing progression axes across cells relies on “pseudotime” analysis, where each cell is assigned a given time value corresponding to its relative state on the main axis or axes of variation. However, due to the destructive nature of scRNA-seq, inferring the trajectory of each cell across timepoints remains experimentally challenging. This motivated the development of TrajectoryNet, a manifold-aware neural ODE network that performs dynamic optimal transport between timepoints^22,23^. TrajectoryNet learns individual trajectories for each cell from the first timepoint (WT) to the last timepoint (*Lep^ob/ob^*) by optimizing for distribution matching between the predicted and true timepoint, as well as energy efficiency, which enables dynamic optimal transport of cells. Additionally, TrajectoryNet is regularized for *unbalanced* dynamic optimal transport through modeling cellular proliferation and death with an auxiliary proliferation network. Together, this ensures TrajectoryNet can learn constrained continuous normalizing flows for each cell that are biologically plausible.

##### TrajectoryNet-based cell-of-origin definition

Each individual trajectory learned by TrajectoryNet defines the trajectory of a given cell in the last timepoint, where TrajectoryNet generates the most likely cells along the path to the first timepoint. Thus, the generated cells at the first timepoint represent the “cells-of-origin” for the learned trajectories. Notably, in addition to distribution matching, TrajectoryNet ensures cellular paths are energy efficient and models biological proliferation and death, meaning that the learned cells-of-origin may be enriched for a particular subset of cells within the first timepoint (in our dataset, cells in the WT population).

##### TrajectoryNet-based granger causality analysis

Using the top 2,278 highly variable genes (defined in *Preprocessing of scRNA-seq data* above), we first computed the mean gene trends over time with TrajectoryNet for two runs averaged. For each transcription factor (TF) in the TRRUST database^72^ and all highly variable genes, we computed the *p*-value of the Granger causality test, or the time-lagged correlation between the TF and the target with lag of 10 (10% of the trajectory), where the *p*-value is computed based on the chi square distribution. We then convert the *p*-value into a Granger causality score, as described previously^23^, by taking the log and adding a sign based on the correlation between two gene trends. For building a regulatory network of genes strongly associated with the obesity progression trajectory, we first identified the top 100 transcription factors with the most regulatory effect (*i.e.*, the highest Granger score across all highly variable genes) and subsetted all TF-target relationships from TRRUST to those TFs. Then, within this subset, we further filtered TF-target pairs to only those with high absolute Granger score (*i.e.*, -log (*p*-value) > 25). We considered this network the full regulatory network. For visualization, we extracted those pairs where both genes are considered “increasing” over the trajectory based on the gene cluster analysis. We visualized the largest connected component, which includes 28 out of 36 genes in the “increasing” subnetwork.

##### Experimental details

We learned cellular trajectories to all cells in the *Lep^ob/ob^*condition for all endocrine cells and to cells assigned to *Lep^ob/ob^*AT 2 for the β cells in our obesity models and in the mSTZ model. TrajectoryNet was run twice in each setting, where enrichment scores (see below) were calculated separately and plotted together, and the mean expression from both runs was computed for visualizing gene trends, calculating gene clusters, and downstream gene set enrichment analysis. One run was used for visualizing individual cellular trajectories. We ran TrajectoryNet on embeddings built from running MAGIC^108^, then PCA (to ensure denoised trajectories as well as invertibility to the gene space), and with the proliferation regularization with alpha = 2 (other parameters default).

#### scMMGAN

##### Overview

Mapping one dataset to another or generating a representation of one dataset in another modality, species, condition, or strain, allows us to build a broader understanding of the broad relationships between cells from different settings toward identifying common gene signatures. scMMGAN is a generative adversarial network architecture for domain adaptation, where the model is trained to preserve relationships between data points and ensure the original data can be reconstructed from the generated output^26^. Unlike batch correction methods, scMMGAN allows us to map β cells from different experimental setups without the obesity progression manifold.

##### Experimental details

For all runs, we used 15,000 training steps, batch size of 128, learning rate of 0.0001, and λ_cycle_ of 1. The correspondence loss for scMMGAN can be flexibly defined. For all mappings, we enforce the preservation of the Pearson correlation of a cell prior to mapping and after mapping to the new domain. This ensures maintenance of the cell’s identity for accurate mappings. For mapping vehicle to WT cells and mSTZ to *Lep^ob/ob^*cells, we ran scMMGAN after MAGIC, then PCA (to match the preprocessing of each condition for AAnet). We used λ_correspondence_ of 10, and scMMGAN was run twice to show consistency of enrichment scores. For mapping the mouse atlas and the human samples, we ran scMMGAN after PCA (for the batch-integrated mouse atlas in 1 run, and for the human datasets individually due to batch effect). Due to the larger differences between these datasets and our obesity progression, versus the mSTZ dataset, we used λ_correspondence_ of 15 to increase preservation of biological identity. The mapping was then visualized using PHATE^106^.

##### Evaluation of integration

To assess integration of batches, we assessed the tradeoff of integration and biological preservation. First, we computed the modified average silhouette width (ASW) of batch, as previously implemented^109^. This metric measures the silhouette of a given batch for each cell type (where we have only one cell type). For the scaled metric we used here, 0 indicates separation of batches, and 1 indicates perfect overlap of batches. To ensure we did not enforce mixing where cells are transcriptionally dissimilar, we also evaluated our integration based on preservation of biological signatures. We thus computed the cosine similarity of each cell before and after mapping (where high cosine similarity is 1 and low cosine similarity is -1). We termed this measure global distortion based on a related metric using cell correlation^110^. A good mapping has both batch integration and cosine similarity near 1, though generally, a tradeoff is necessary due to large technical variation between datasets. In particular, we expected some transformation of cellular gene expression to ensure mapping. Thus, we aimed for mild to moderate global distortion (cosine similarity = 0.2-0.6) and high batch integration (integration > 0.6) in order to ensure the nearest cells in the obesity progression were truly near in the embedding space.

For vehicle to WT mapping, scMMGAN balanced batch integration (mean Batch ASW=0.91) and biological preservation (mean cosine similarity before versus after correction=0.69) (**Table S4A**). For mSTZ to *Lep^ob/ob^*, scMMGAN also observed high integration performance (mean Batch ASW=0.64, mean cosine similarity=0.71; **Table S4B**). For the mouse integrated atlas, after scMMGAN alignment, the datasets showed high batch integration (Batch ASW=0.98) and mild global distortion across all cells (cosine similarity=0.58 before and after alignment), with no distortion specific to a particular archetype, developmental stage, disease state, or dataset (**Table S4C**). For human samples, scMMGAN showed high batch integration (Batch ASW=0.97) and moderate global distortion across all cells (cosine similarity=0.35 before and after alignment), common when mapping between datasets with substantial differences, including species-specific differences, with no distortion specific to a particular archetype, sex, or disease state (**Table S4D**).

#### AAnet

##### Overview

Defining cell states via cellular clustering remains a useful tool for single-cell analysis but proves insufficient for characterizing heterogeneity within cell types. Indeed, β cell heterogeneity is well appreciated in the field, but cell states are highly plastic and require an alternative approach to model this plasticity. AAnet^24,25^ is a latent-space archetypal analysis approach that learns cellular “archetypes”, or extreme states, and defines each cell with respect to the extreme states. This allows us to identify cells highly specific to a given state.

##### Archetypal commitment

Archetypal commitment is captured in the AAnet latent space, where each cell is defined by its affinity to each archetype (a vector of size number of archetypes, where the entries sum to 1 and higher value implies higher commitment). Thus, we determine cells that have commitment over 0.5 for a given archetype as “committed” to that archetype^25^.

##### Archetypal assignment for mapped cells

For the vehicle and mSTZ analysis^56^, we learned mappings directly onto the embedding space used for training the WT AAnet and *Lep^ob/ob^* AAnet models, respectively. Thus, for these datasets, we used the trained AAnet models to encode archetypal assignments for cells. For other mappings, including with TrajectoryNet and with scMMGAN on the full trajectory of cells, we assigned each cell the archetype of its closest neighbor in the reference dataset.

##### Experimental details

We ran AAnet with default parameters on each condition embedding separately, defined by running MAGIC followed by PCA (again to enable denoising and invertibility to the gene space). To determine the number of archetypes for each condition^24^, we first generated a training and test set, where the training test was generated by density subsampling 80% of the dataset, and the test set was the held-out group. We then trained AAnet on the training set 3 times for each number of archetypes between 2 and 10. For each trained model, we calculated the mean squared error (MSE) of the input versus reconstructed cells for the held-out test set. We plotted the MSE and chose the elbow point as the number of archetypes (**Figure S5A**).

#### DiffusionEMD

##### Overview

Patient manifolds are multi-scale manifolds, where one level characterizes relationships between cells, and a higher level characterizes relationships between patients^111^. To compare single-cell datasets between patients, such as to compare cell type proportions, gene trends, or clinical variables across patient mappings, Diffusion Earth Mover’s Distance (DiffusionEMD)^27^ models each single-cell data set as a distribution supported on a common cell-cell graph enabling comparison via optimal transport.

##### Experimental details

We ran DiffusionEMD with default parameters to embed human patient samples after their mapping to the obesity progression with scMMGAN.

###### Enrichment score calculation

To compute the enrichment of a given label in a mapped population of cells, we annotated each mapped cell using the label of the nearest “real” cell in the reference dataset. We then calculated the proportion of each class in the mapped population. This alone does not sufficiently characterize the mapping of a population due to large class imbalances between labels. Therefore, we calculated the ratio between the mapped label proportions and the label proportions in the reference dataset. We term this the “enrichment score”. For each class, if the ratio is greater than 1, the class is “enriched” in the mapped population. If the ratio is less than 1, the class is enriched in the reference population. If the ratio is equal to 1, the class is equally represented in the reference and the mapped population. For all endocrine cells (**Figure S3**), we calculated the enrichment of each cell type for cells-of-origin to *Cck*+ *Lep^ob/ob^* β cells and for cells-of-origin to *Cck*+ *Lep^ob/ob^* polyhormonal cells versus the distribution of cell types in the WT population. Within β cells (**Figure 3**), we calculated the enrichment of each WT archetype for cells-of-origin versus the distribution of archetypes in the WT population. We also calculated the enrichment of each archetype for each timepoint versus the distribution of all archetypes to learn the trajectory of archetypes from WT AT 4 to *Lep^ob/ob^* AT 2. For the mSTZ dataset (**Figure 5**), we calculated the enrichment of each WT archetype for the β2 population versus the distribution of archetypes in the reference WT population. We also calculated the enrichment of each *Lep^ob/ob^*archetype for the *Cck*-hi population versus the distribution of *Lep^ob/ob^* archetypes in the reference. Finally, we calculated the enrichment of β1 versus β2 for mapped cells-of-origin from TrajectoryNet to *Lep^ob/ob^*AT 2-annotated cells.

###### Differential expression

For differential gene expression of archetypes, we compared cells committed to each archetype versus all other cells and performed a Wilcoxon rank sum test, where differentially expressed genes have BH-adjusted *p*-value (*q*-value) < 0.05 and log fold-change > 0.0.

###### Gene clusters

We clustered the 2,278 highly variable genes based on TrajectoryNet-inferred transcriptional changes from all WT to *Lep^ob/ob^* AT 2. We computed the mean expression over all trajectories from two TrajectoryNet runs and clustered each gene’s association with continuous time. Genes were clustered using *k* means clustering with the number of clusters set to two.

###### Gene set enrichment analysis

Gene set enrichment analysis was performed using GSEApy^65^ and Enrichr^66^, using the BioPlanet 2019, Panther 2016, KEGG 2019, Reactome 2022, and GO Biological Process 2023 databases. Gene sets with BH-adjusted *p*-value (*q*-value) < 0.05 were retained. The hypergeometric test was used for determining statistical significance of overlapping gene sets.

###### CUT&RUN analysis

Adapter sequences from paired-end sequencing reads were trimmed with Trimmomatic and quality control was performed using FastQC. Bowtie2 version 2.4.2 was used to align reads to the GRCm38 genome using the –very-sensitive flag and allowing for one mismatch. SAMtools version 1.16 was used to remove unmapped reads, secondary alignment reads, and reads with a Phred score of <30. Macs2 version 2.2.7.1 was used to call narrow peaks with default peak-calling settings. H3K4me3 and cJun CUT&RUN peaks were called against IgG negative controls. IDR was used to determine significant peaks between the two biological replicates. Deeptools version 3.5.1 was used for data visualization. To visualize average CUT&RUN signal per sample, Deeptools bamCoverage was used to normalize bigWig signal based on E. coli spike-in control DNA. A scaling factor was calculated based on the percentage of reads mapped to the E. coli K12, MG1655 reference genome. The following additional bamCoverage flags were used: --extendReads --ignoreDuplicates --samFlagInclude 64. Differences in read coverage between samples were normalized based on CPM. In all plots, the mean normalized bigWig signal between two biological replicates is plotted. Peak intersections were performed using BEDTools v2.30. Enriched motifs were identified using MEME v5.4.1 meme-chip.

###### Analysis of publicly available ChIP-seq and ATAC-seq data

The following published datasets from ENCODE^75,76^ were used: H3K27ac ChIP-seq from mouse cortex (ENCSR491RBV); H3K27ac ChIP-seq from mouse hippocampus (ENCSR527FRO); DNase-seq from mouse hippocampus (ENCSR544UYQ); and CTCF ChIP-seq from human adult pancreas tissue (ENCSR774PGN). H3K27ac ChIP-seq and ATAC-seq from human islets were obtained from Miguel-Escalada et al.^77^. Loop anchor and Hi-C data from human islets were obtained from Greenwald et al.^78^. For all published and publicly available datasets used, BED and/or bigWig files were converted to the mm10 genome or hg19 genome for mouse and human sequencing datasets, respectively. Peaks and bigWig signal were visualized with IGV. Hi-C data was visualized with the WashU Epigenome Browser^112–115^.

## RESOURCE AVAILABILITY

### Lead Contact

Requests for further information and resources should be directed to and will be fulfilled by the Lead Contact, Mandar Deepak Muzumdar.

### Materials Availability

All unique/stable reagents generated in this study are available from the Lead Contact with a completed Materials Transfer Agreement.

### Data and code availability

Newly generated scRNA-seq data and CUT&RUN sequencing data were deposited into the Gene Expression Omnibus (GEO) under accession numbers GSE279485 and GSE280947, respectively. Accession numbers for reanalyzed data are listed above. All code used in sequencing analyses is available on GitHub (https://github.com/KrishnaswamyLab/Beta-Cell-Driven-PDAC). All other raw and analyzed data is available upon request.

## SUPPLEMENTAL INFORMATION

**Figure S1.**
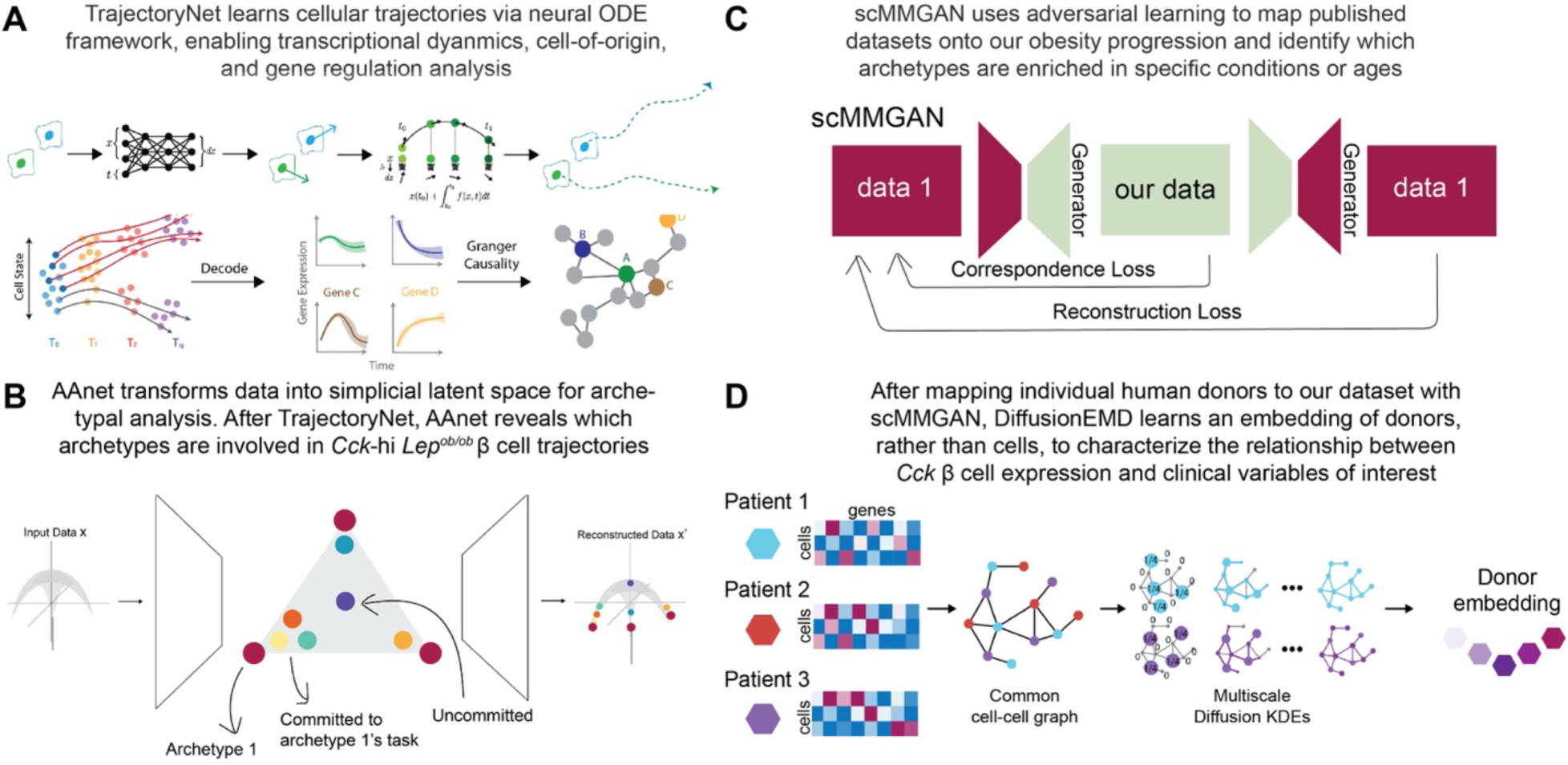
Overview of computational methods used in the study, Related to Figures 2, 3, 5, and 6. (A) TrajectoryNet overview. TrajectoryNet performs neural ODE-based optimal transport, enabling transcriptional dynamics, cell-of-origin, and gene regulation analysis. (B) AAnet overview. AAnet transforms data into latent representation shaped like a simplex for performing latent-space archetypal analysis. Combined with TrajectoryNet, AAnet characterizes archetype-of-origin and archetypes involved in Cck-hi *Lep^ob/ob^* β cell trajectories. (C) scMMGAN overview. scMMGAN is a generative adversarial network, using adversarial training to map non-diabetic and type II diabetic (T2D) datasets onto our obesity progression. Combined with AAnet, scMMGAN identifies which archetypes are enriched in the context of different stressors or developmental stages. Combined with TrajectoryNet, scMMGAN assesses if the mapped cells show the same cellular dynamics as the obesity progression. (D) DiffusionEMD overview. Mapping scRNA-seq datasets from human donors with scMMGAN shows similarity between cells from human donors versus our dataset. DiffusionEMD builds an embedding of donors based on similarity of cellular populations between donors, which enables comparison of donor-level attributes (e.g. age, sex, T2D status) and enrichment along the obesity progression.

**Figure S2.**
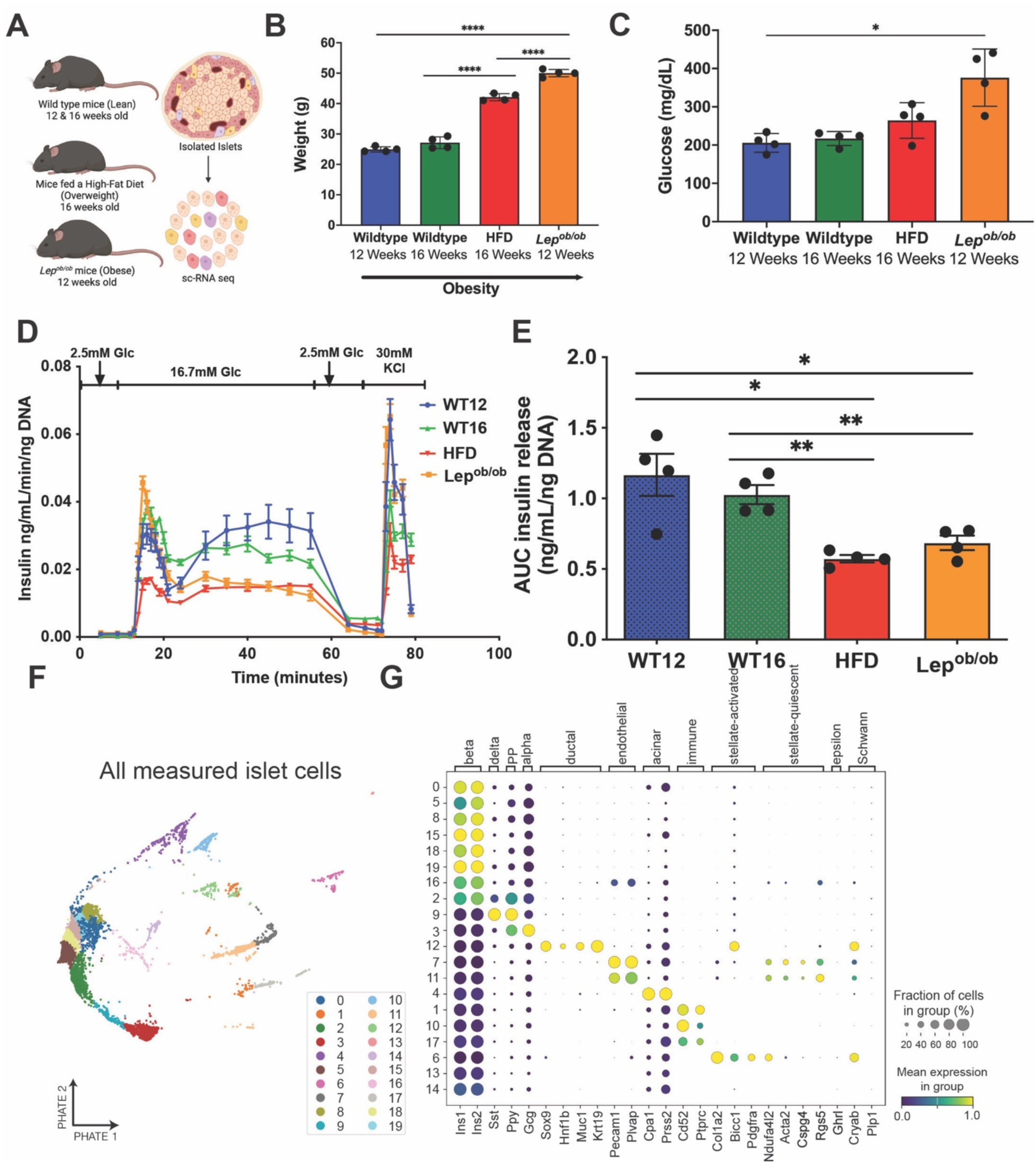
TrajectoryNet enables the identification of the cell-of-origin for aberrant β cell expression of cholecystokinin, Related to Figures 2 and 3. (A) Overview of scRNA-seq experiment on mouse islets from congenic obesity models (high-fat diet (HFD)-fed for 10 weeks, *Lep^ob/ob^*) and age-matched wild-type (WT) controls. Generated using Biorender.com. (B) Mice weight (mean+/-s.e.m., n=4 mice per group) at time of harvest (age listed for each group). ****p<0.0001, Welch’s t-test. (C) Random glucose levels (mean+/-s.e.m., n=4 mice per group) at time of harvest (age listed for each group). *p<0.05, Welch’s t-test. (D) Glucose stimulated insulin secretion (GSIS) of islets isolated from congenic WT (12 and 16 weeks of age (WT12, WT16)), 16-week-old mice fed HFD-fed mice, and 12-week-old *Lep^ob/ob^* mice. Data from WT12 and *Lep^ob/ob^*were reanalyzed from our prior work^18^. (E) The area under the curve (AUC; mean+/-s.e.m., n=4 replicates per group) for GSIS in (**D**). *p<0.05, **p<0.01, Welch’s t test. (F) PHATE embedding of exocrine and endocrine islet cells, colored by *k*-means cluster annotation. (G) Dot plot of cluster expression for marker genes pertaining to islet populations. Clusters 0, 5, 8, 15, 16, 18, and 19, were annotated as β cells; 2 as polyhormonal (insulin and another hormone); 9 as δ and PP cells; 3 as α cells; 12 as duct cells; 7 and 11 as endothelial cells; 4 as acinar cells; 1, 10, 17 as immune cells; 6 as stellate-activated cells; and 13, 14 as low-quality clusters.

**Figure S3.**
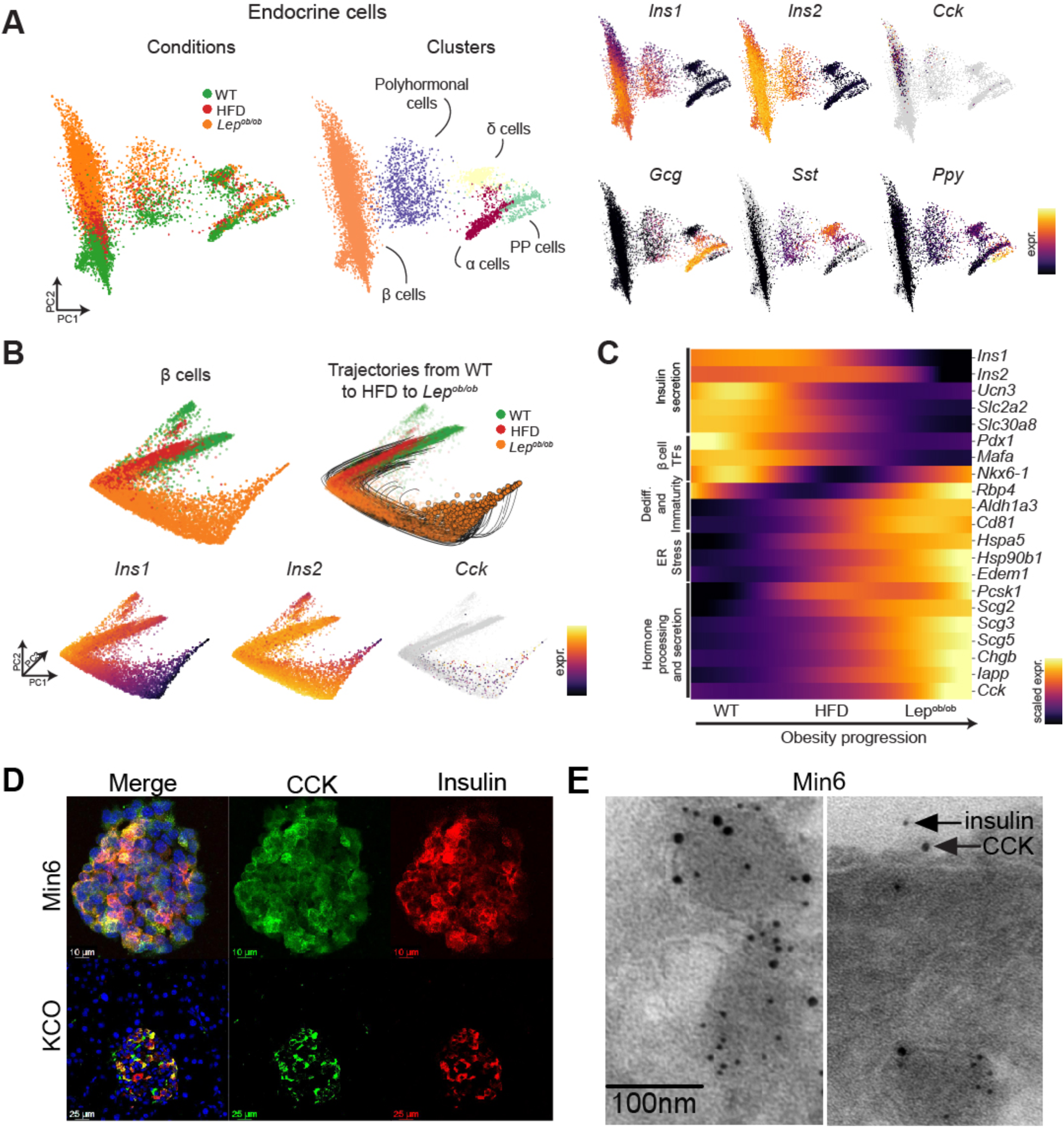
Dynamic changes in β cell expression of *insulin* and *Cck* with obesity, Related to Figures 3 and 4. (A) Combined embedding of endocrine cells from three conditions (WT, HFD, *Lep^ob/ob^*) and annotated clusters based on marker genes. (B) β cell embedding colored by condition (top) and insulin (*Ins1*, *Ins2*) and *Cck* expression. (C) Mean gene expression across all cells along the obesity progression axis for key marker genes. Visualized trajectories to *Cck*-hi cells on this axis are shown in (**B**). (D) Representative co-immunofluorescence images of mouse insulinoma cells (Min6) and *KCO* mice displaying nuclei (DAPI, blue), CCK (green), and insulin (red). (E) Representative co-immunoelectron microscopy images of Min6 cells labeled with insulin (5 nm dots) and CCK (10 nm dots). Arrows denote insulin and CCK at the plasma membrane, indicating possible active co-secretion.

**Figure S4.**
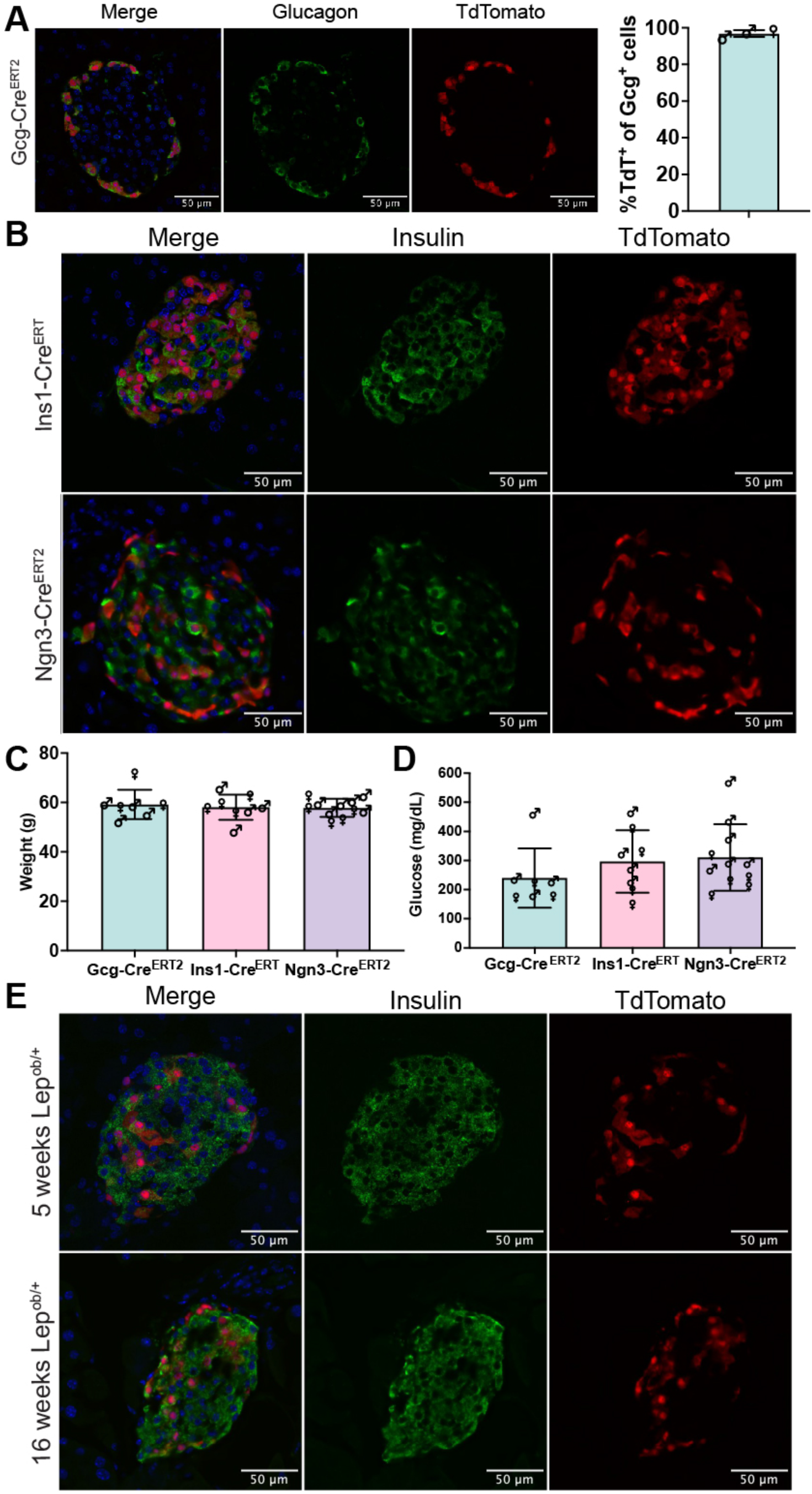
Validation of CreER lines for *in vivo* lineage tracing, Related to Figure 4. (A) Representative images of Gcg immunofluorescence and lineage-traced TdTomato+ cells in 5-week-old *Gcg-Cre^ERT^*^2^*; Lep^ob/ob^; Rosa26^LSL-TdTomato^* mice administered tamoxifen at 4 weeks of age. DAPI labels nuclei blue. Average percentage (mean+/-s.e.m., n=3 mice, sex designated by symbols) of Gcg+ cells labeled with TdTomato in mice is shown showing near complete overlap. (B) Representative images of insulin immunofluorescence and lineage-traced TdTomato+ cells in 5-week-old *Ins1-Cre^ERT^; Lep^ob/ob^; Rosa26^LSL-TdTomato^* and *Ngn3-Cre^ERT^*^2^*; Lep^ob/ob^; Rosa26^LSL-TdTomato^* mice administered tamoxifen at 4 weeks of age. DAPI labels nuclei blue. (C) Final weights of 16-week-old mice (*Lep^ob/ob^*) for each mouse line (mean+/-s.e.m., n=7-10 mice per group, sex is designated by symbols) in Figure 4B. Data are not significantly different between groups, Welch’s t-test. (D) Final random glucose levels of 16-week-old mice (*Lep^ob/ob^*) for each mouse line (mean+/- s.e.m., n=7-10 mice per group, sex is designated by symbols) in Figure 4B. Data are not significantly different between groups, Welch’s t-test. (E) Representative images of insulin immunofluorescence and lineage-traced TdTomato+ cells in 5-week-old and 16-week-old lean *Ngn3-Cre^ERT^*^2^*; Lep^ob/+^; Rosa26^LSL-TdTomato^* mice administered tamoxifen at 4 weeks of age. DAPI labels nuclei blue.

**Figure S5.**
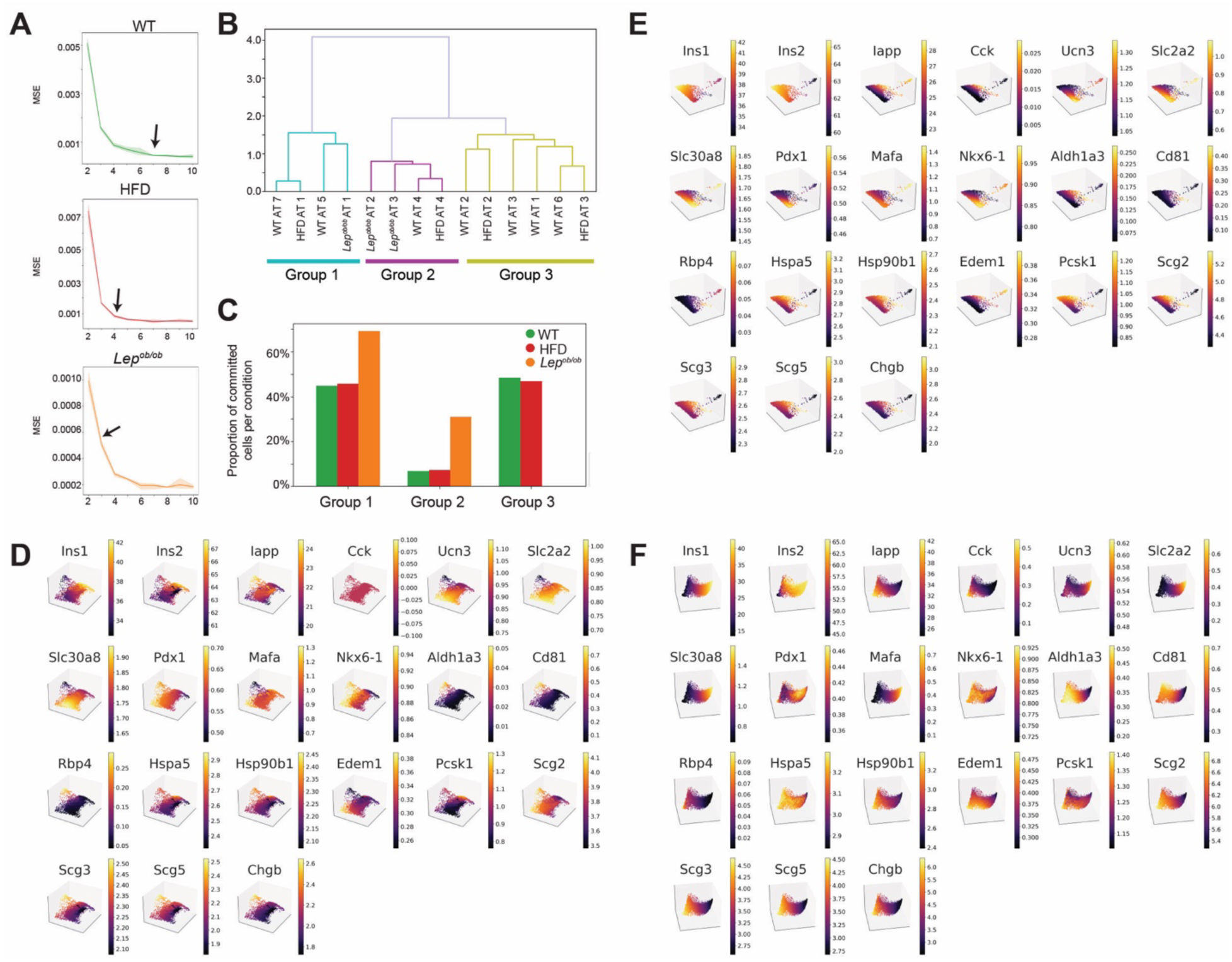
Analysis of β cell heterogeneity within WT, HFD, and *Lep^ob/ob^* embeddings, Related to Figures 3 and 5. (A) AAnet-inferred number of archetypes, computed based on elbow point of reconstruction error for held-out test set over number of archetypes from [2,10], for three runs per archetype. (B) Cosine similarity between archetypes defined across the entire transcriptome reveals three groups of archetypes shared across WT, HFD, and *Lep^ob/ob^* settings. (C) Proportion of committed cells from each condition per archetypal group. (D) WT embedding of archetypes colored by marker genes. (E) HFD embedding of archetypes colored by marker genes. (F) *Lep^ob/ob^* embedding of archetypes colored by marker genes.

**Figure S6.**
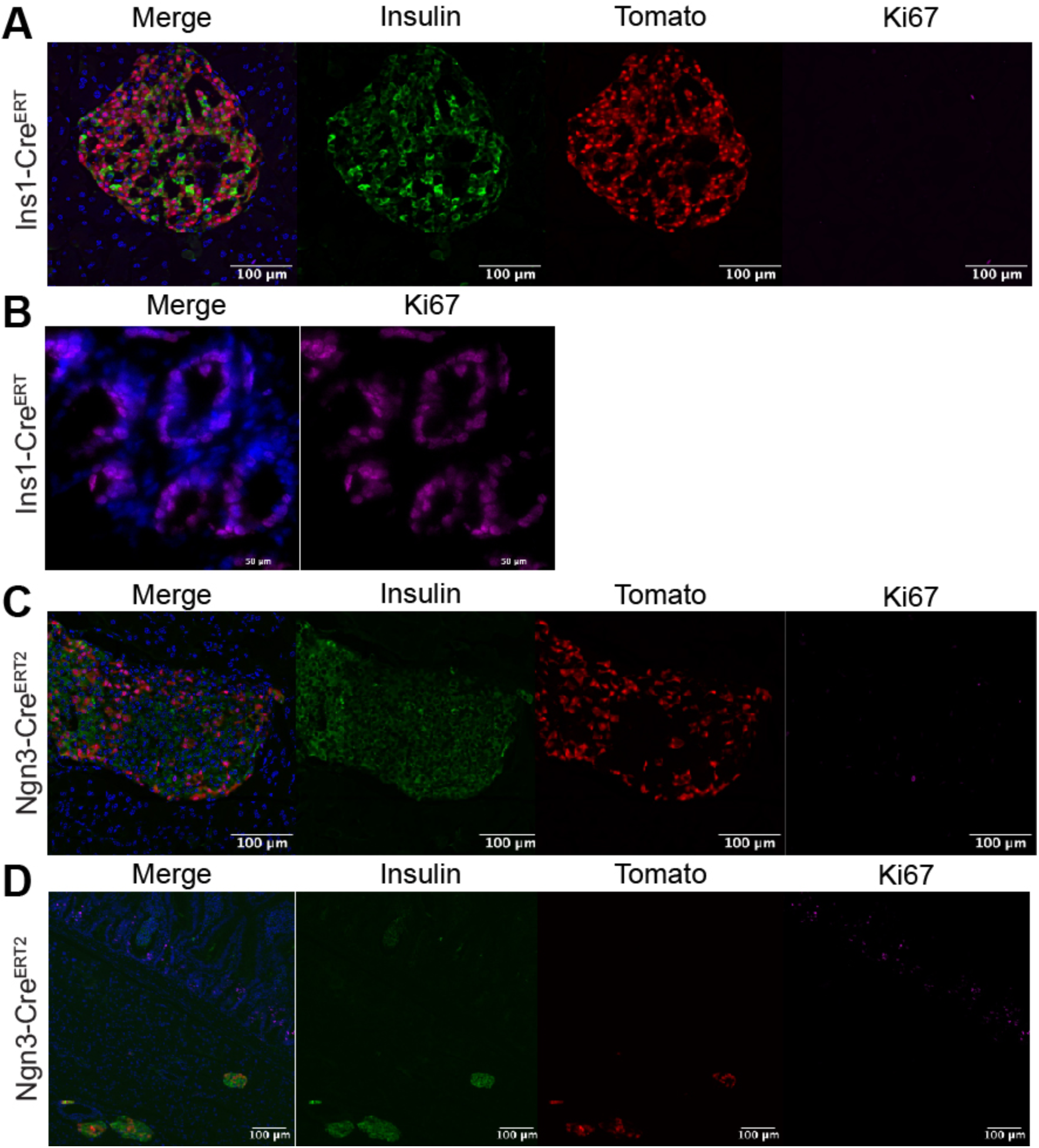
Absence of actively proliferating β cells at 16 weeks in *Lep^ob/ob^* mice, Related to Figure 4. (A) Co-immunofluorescence for insulin and Ki67 of islets from 16-week-old *Ins1-Cre^ERT^; Lep^ob/ob^; Rosa26^LSL-TdTomato^*mice shows no Ki67+ cells within the islet (DAPI: blue, insulin: green, TdTomato: red, Ki67: magenta). (B) Immunofluorescence for Ki67 (magenta) in small intestine from 16-week-old *Ins1-Cre^ERT^; Lep^ob/ob^; Rosa26^LSL-TdTomato^* mice demonstrates Ki67+ cells in the crypts as a positive control. DAPI labels nuclei blue. (C) Co-immunofluorescence for insulin and Ki67 of islets from 16-week-old *Ngn3-Cre^ERT^*^2^*; Lep^ob/ob^; Rosa26^LSL-TdTomato^*mice shows no Ki67+ cells within the islet (DAPI: blue, insulin: green, TdTomato: red, Ki67: magenta). (D) Co-immunofluorescence for insulin and Ki67 of islets from 16-week-old *Ngn3-Cre^ERT^*^2^*; Lep^ob/ob^; Rosa26^LSL-TdTomato^*mice shows no Ki67+ cells within the islet (DAPI: blue, insulin: green, TdTomato: red, Ki67: magenta) but the presence of Ki67+ cells in the intestinal crypts (top).

**Figure S7.**
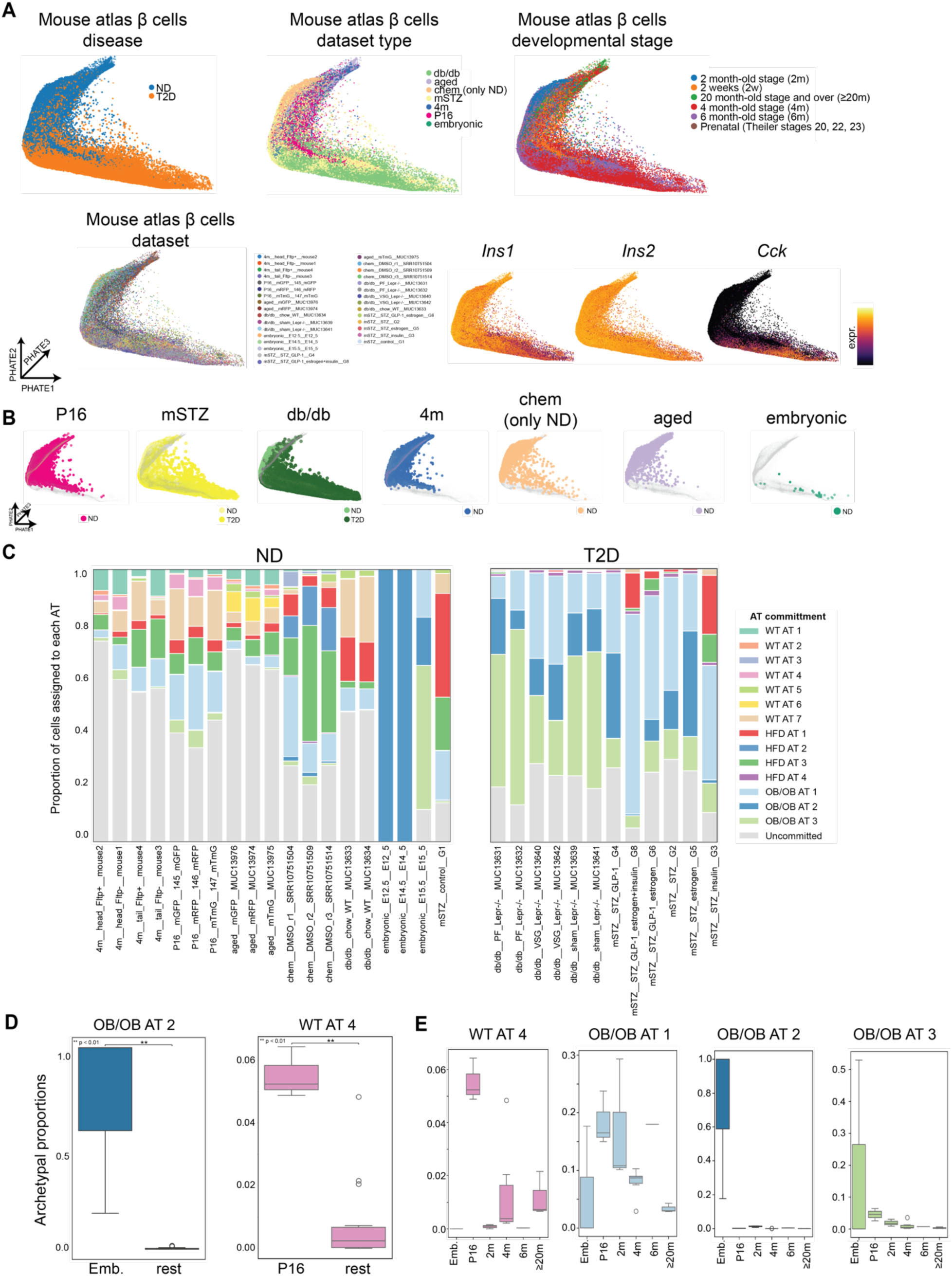
Comparison of obesity progression to β cells from non-diabetic (ND) and type II diabetes (T2D) mouse models, Related to Figure 6. (A) Embedding of mouse atlas β cells mapped onto the obesity progression with scMMGAN and colored by disease annotation, dataset type, development stage, specific dataset, and marker genes. (B) Individual embeddings of each mouse dataset mapped to the obesity progression. (C) Proportion of each archetype for each ND and T2D dataset. (D) Archetypal proportion for Embryonic and P16 samples versus other samples. Box plots display 25^th^, 50^th^, and 75^th^ percentiles +/- 1.5 interquartile range (IQR). **p<0.01, Wilcoxon rank sum test. (E) Archetypal proportion for WT AT 4, *Lep^ob/ob^* AT 1, *Lep^ob/ob^* AT 2, and *Lep^ob/ob^*AT 3 for ND datasets over age (embryonic to aged mice). Box plots display 25^th^, 50^th^, and 75^th^ percentiles +/- 1.5 interquartile range (IQR).

## SUPPLEMENTARY TABLES

**Table S1, related to Figure 3**. Wilcoxon rank sum test results for all genes (log fold-change and Benjamini-Hochberg (BH)-adjusted *p*-value) for each archetype versus rest within each condition (WT, HFD, and *Lep^ob/ob^*(ob.ob)).

**Table S2, related to Figure 5**. Enriched (*q* < 0.05) genes from gene set enrichment analysis with BioPlanet, KEGG, Reactome, and GO Biological Process for genes that (A) decrease and (B) increase in expression along the obesity progression axis.

**Table S3, related to Figure 5**. Gene sets and associated genes enriched with respect to each human stressor and genes increasing over obesity progression.

**Table S4, related to Figures 5** and **6**. scMMGAN evaluation of mapping with batch ASW and global distortion (cosine similarity before and after alignment) for (A) vehicle cells (from mSTZ experiment) to WT cells, (B) mSTZ-treated cells to *Lep^ob/ob^* cells, (C) mouse atlas cells to WT, HFD, and *Lep^ob/ob^* cells, and (D) human ND and T2D donor cells to WT, HFD, and *Lep^ob/ob^* cells.

**Table S5, related to Figure 7**. Gene regulatory network, with subnetwork of genes identified as increasing over the obesity trajectory annotated and visualized in **Figure 7B**.

**Table S6, related to Figure 7**. Enriched (*q* < 0.05) genes from gene set enrichment analysis with BioPlanet, KEGG, Reactome, and GO Biological Process for subnetwork of genes identified as increasing over the obesity trajectory.

## REFERENCES

1. Mastracci, T.L., Apte, M., Amundadottir, L.T., Alvarsson, A., Artandi, S., Bellin, M.D., Bernal-Mizrachi, E., Caicedo, A., Campbell-Thompson, M., Cruz-Monserrate, Z., et al. (2022). Integrated Physiology of the Exocrine and Endocrine Compartments in Pancreatic Diseases: Workshop Proceedings. Pancreas 51, 1061–1073. 10.1097/mpa.0000000000002170.

2. Overton, D.L., and Mastracci, T.L. (2022). Exocrine-Endocrine Crosstalk: The Influence of Pancreatic Cellular Communications on Organ Growth, Function and Disease. Front Endocrinol (Lausanne) 13, 904004. 10.3389/fendo.2022.904004.

3. Raeder, H., Johansson, S., Holm, P.I., Haldorsen, I.S., Mas, E., Sbarra, V., Nermoen, I., Eide, S.A., Grevle, L., Bjørkhaug, L., et al. (2006). Mutations in the CEL VNTR cause a syndrome of diabetes and pancreatic exocrine dysfunction. Nat Genet 38, 54–62. 10.1038/ng1708.

4. Kahraman, S., Dirice, E., Basile, G., Diegisser, D., Alam, J., Johansson, B.B., Gupta, M.K., Hu, J., Huang, L., Soh, C.L., et al. (2022). Abnormal exocrine-endocrine cell cross-talk promotes beta-cell dysfunction and loss in MODY8. Nat Metab 4, 76–89. 10.1038/s42255-021-00516-2.

5. Xu, M., Jung, X., Hines, O.J., Eibl, G., and Chen, Y. (2018). Obesity and Pancreatic Cancer: Overview of Epidemiology and Potential Prevention by Weight Loss. Pancreas 47, 158–162. 10.1097/MPA.0000000000000974.

6. Cascetta, P., Cavaliere, A., Piro, G., Torroni, L., Santoro, R., Tortora, G., Melisi, D., and Carbone, C. (2018). Pancreatic Cancer and Obesity: Molecular Mechanisms of Cell Transformation and Chemoresistance. Int J Mol Sci 19 10.3390/ijms19113331.

7. Yuan, C., Bao, Y., Wu, C., Kraft, P., Ogino, S., Ng, K., Qian, Z.R., Rubinson, D.A., Stampfer, M.J., Giovannucci, E.L., and Wolpin, B.M. (2013). Prediagnostic body mass index and pancreatic cancer survival. J Clin Oncol 31, 4229–4234. 10.1200/JCO.2013.51.7532.

8. Tan, J., You, Y., Guo, F., Xu, J., Dai, H., and Bie, P. (2017). Association of elevated risk of pancreatic cancer in diabetic patients: A systematic review and meta-analysis. Oncol Lett 13, 1247–1255. 10.3892/ol.2017.5586.

9. Kasenda, B., Bass, A., Koeberle, D., Pestalozzi, B., Borner, M., Herrmann, R., Jost, L., Lohri, A., and Hess, V. (2014). Survival in overweight patients with advanced pancreatic carcinoma: a multicentre cohort study. BMC Cancer 14, 728. 10.1186/1471-2407-14-728.

10. Sharma, A., and Chari, S.T. (2018). Pancreatic Cancer and Diabetes Mellitus. Curr Treat Options Gastroenterol 16, 466–478. 10.1007/s11938-018-0197-8.

11. Storz, P., and Crawford, H.C. (2020). Carcinogenesis of Pancreatic Ductal Adenocarcinoma. Gastroenterology 158, 2072–2081. 10.1053/j.gastro.2020.02.059.

12. Kopp, J.L., von Figura, G., Mayes, E., Liu, F.F., Dubois, C.L., Morris, J.P.t., Pan, F.C., Akiyama, H., Wright, C.V., Jensen, K., et al. (2012). Identification of Sox9-dependent acinar-to-ductal reprogramming as the principal mechanism for initiation of pancreatic ductal adenocarcinoma. Cancer Cell 22, 737–750. 10.1016/j.ccr.2012.10.025.

13. Zhang, A.M.Y., Magrill, J., de Winter, T.J.J., Hu, X., Skovso, S., Schaeffer, D.F., Kopp, J.L., and Johnson, J.D. (2019). Endogenous Hyperinsulinemia Contributes to Pancreatic Cancer Development. Cell Metab 30, 403–404. 10.1016/j.cmet.2019.07.003.

14. Zhang, A.M.Y., Chu, K.H., Daly, B.F., Ruiter, T., Dou, Y., Yang, J.C.C., de Winter, T.J.J., Chhuor, J., Wang, S., Flibotte, S., et al. (2022). Effects of hyperinsulinemia on pancreatic cancer development and the immune microenvironment revealed through single-cell transcriptomics. Cancer Metab 10, 5. 10.1186/s40170-022-00282-z.

15. Zhang, A.M.Y., Xia, Y.H., Lin, J.S.H., Chu, K.H., Wang, W.C.K., Ruiter, T.J.J., Yang, J.C.C., Chen, N., Chhuor, J., Patil, S., et al. (2023). Hyperinsulinemia acts via acinar insulin receptors to initiate pancreatic cancer by increasing digestive enzyme production and inflammation. Cell Metab 35, 2119–2135.e2115. 10.1016/j.cmet.2023.10.003.

16. Storz, P. (2017). Acinar cell plasticity and development of pancreatic ductal adenocarcinoma. Nat Rev Gastroenterol Hepatol 14, 296–304. 10.1038/nrgastro.2017.12.

17. Friedman, J.M. (2019). Leptin and the endocrine control of energy balance. Nature Metabolism. 10.1038/s42255-019-0095-y.

18. Chung, K.M., Singh, J., Lawres, L., Dorans, K.J., Garcia, C., Burkhardt, D.B., Robbins, R., Bhutkar, A., Cardone, R., Zhao, X., et al. (2020). Endocrine-Exocrine Signaling Drives Obesity-Associated Pancreatic Ductal Adenocarcinoma. Cell 181, 832–847 e818. 10.1016/j.cell.2020.03.062.

19. Ardito, C.M., Gruner, B.M., Takeuchi, K.K., Lubeseder-Martellato, C., Teichmann, N., Mazur, P.K., Delgiorno, K.E., Carpenter, E.S., Halbrook, C.J., Hall, J.C., et al. (2012). EGF receptor is required for KRAS-induced pancreatic tumorigenesis. Cancer Cell 22, 304–317. 10.1016/j.ccr.2012.07.024.

20. Williams, J.A. (2019). Cholecystokinin (CCK) Regulation of Pancreatic Acinar Cells: Physiological Actions and Signal Transduction Mechanisms. In Comprehensive Physiology, pp. 535–564. 10.1002/cphy.c180014.

21. Guerra, C., Schuhmacher, A.J., Canamero, M., Grippo, P.J., Verdaguer, L., Perez-Gallego, L., Dubus, P., Sandgren, E.P., and Barbacid, M. (2007). Chronic pancreatitis is essential for induction of pancreatic ductal adenocarcinoma by K-Ras oncogenes in adult mice. Cancer Cell 11, 291–302. 10.1016/j.ccr.2007.01.012.

22. Tong, A., Huang, J., Wolf, G., van Dijk, D., and Krishnaswamy, S. (2020). TrajectoryNet: A Dynamic Optimal Transport Network for Modeling Cellular Dynamics. Proc Mach Learn Res 119, 9526–9536.

23. Tong, A., Kuchroo, M., Gupta, S., Venkat, A., Juan, B.P.S., Rangel, L., Zhu, B., Lock, J.G., Chaffer, C.L., and Krishnaswamy, S. (2023). Learning transcriptional and regulatory dynamics driving cancer cell plasticity using neural ODE-based optimal transport. bioRxiv, 2023.2003.2028.534644. 10.1101/2023.03.28.534644.

24. van Dijk, D., Burkhardt, D.B., Amodio, M., Tong, A., Wolf, G., and Krishnaswamy, S. (2019). Finding archetypal spaces using neural networks. arXiv 1901.09078v2. https://arxiv.org/abs/1901.09078v2.

25. Venkat, A., Youlten, S.E., Juan, B.P.S., Purcell, C., Amodio, M., Burkhardt, D.B., Benz, A., Holst, J., McCool, C., Mollbrink, A., et al. (2024). AAnet resolves a continuum of spatially-localized cell states to unveil tumor complexity. bioRxiv, 2024.2005.2011.593705. 10.1101/2024.05.11.593705.

26. Amodio, M., Youlten, S.E., Venkat, A., San Juan, B.P., Chaffer, C.L., and Krishnaswamy, S. (2022). Single-cell multi-modal GAN reveals spatial patterns in single-cell data from triple-negative breast cancer. Patterns (N Y) 3, 100577. 10.1016/j.patter.2022.100577.

27. Tong, A., Huguet, G., Natik, A., MacDonald, K., Kuchroo, M., Coifman, R., Wolf, G., and Krishnaswamy, S. (2021). Diffusion Earth Mover’s Distance and Distribution Embeddings. ArXiv.

28. Hingorani, S.R., Wang, L., Multani, A.S., Combs, C., Deramaudt, T.B., Hruban, R.H., Rustgi, A.K., Chang, S., and Tuveson, D.A. (2005). Trp53R172H and KrasG12D cooperate to promote chromosomal instability and widely metastatic pancreatic ductal adenocarcinoma in mice. Cancer Cell 7, 469–483. 10.1016/j.ccr.2005.04.023.

29. Wang, J., Takeuchi, T., Tanaka, S., Kubo, S.K., Kayo, T., Lu, D., Takata, K., Koizumi, A., and Izumi, T. (1999). A mutation in the insulin 2 gene induces diabetes with severe pancreatic beta-cell dysfunction in the Mody mouse. J Clin Invest 103, 27–37. 10.1172/JCI4431.

30. Khasawneh, J., Schulz, M.D., Walch, A., Rozman, J., Hrabe de Angelis, M., Klingenspor, M., Buck, A., Schwaiger, M., Saur, D., Schmid, R.M., et al. (2009). Inflammation and mitochondrial fatty acid beta-oxidation link obesity to early tumor promotion. Proc Natl Acad Sci U S A 106, 3354–3359. 10.1073/pnas.0802864106.

31. Philip, B., Roland, C.L., Daniluk, J., Liu, Y., Chatterjee, D., Gomez, S.B., Ji, B., Huang, H., Wang, H., Fleming, J.B., et al. (2013). A high-fat diet activates oncogenic Kras and COX2 to induce development of pancreatic ductal adenocarcinoma in mice. Gastroenterology 145, 1449–1458. 10.1053/j.gastro.2013.08.018.

32. Chang, H.-H., Moro, A., Takakura, K., Su, H.-Y., Mo, A., Nakanishi, M., Waldron, R.T., French, S.W., Dawson, D.W., Hines, O.J., et al. (2017). Incidence of pancreatic cancer is dramatically increased by a high fat, high calorie diet in KrasG12D mice. Plos One 12. 10.1371/journal.pone.0184455.

33. Incio, J., Liu, H., Suboj, P., Chin, S.M., Chen, I.X., Pinter, M., Ng, M.R., Nia, H.T., Grahovac, J., Kao, S., et al. (2016). Obesity-Induced Inflammation and Desmoplasia Promote Pancreatic Cancer Progression and Resistance to Chemotherapy. Cancer Discovery 6, 852–869. 10.1158/2159-8290.Cd-15-1177.

34. Zyromski, N.J., Mathur, A., Pitt, H.A., Wade, T.E., Wang, S., Nakshatri, P., Swartz-Basile, D.A., and Nakshatri, H. (2009). Obesity potentiates the growth and dissemination of pancreatic cancer. Surgery 146, 258–263. 10.1016/j.surg.2009.02.024.

35. Hrovatin, K., Bastidas-Ponce, A., Bakhti, M., Zappia, L., Buttner, M., Salinno, C., Sterr, M., Bottcher, A., Migliorini, A., Lickert, H., and Theis, F.J. (2023). Delineating mouse beta-cell identity during lifetime and in diabetes with a single cell atlas. Nat Metab 5, 1615–1637. 10.1038/s42255-023-00876-x.

36. Chu, C.M.J., Modi, H., Ellis, C., Krentz, N.A.J., Skovsø, S., Zhao, Y.B., Cen, H., Noursadeghi, N., Panzhinskiy, E., Hu, X., et al. (2022). Dynamic Ins2 Gene Activity Defines β-Cell Maturity States. Diabetes 71, 2612–2631. 10.2337/db21-1065.

37. Szabat, M., Pourghaderi, P., Soukhatcheva, G., Verchere, C.B., Warnock, G.L., Piret, J.M., and Johnson, J.D. (2011). Kinetics and genomic profiling of adult human and mouse β-cell maturation. Islets 3, 175–187. 10.4161/isl.3.4.15881.

38. Luecken, M.D., and Theis, F.J. (2019). Current best practices in single-cell RNA-seq analysis: a tutorial. Mol Syst Biol 15, e8746. 10.15252/msb.20188746.

39. Neelankal John, A., Ram, R., and Jiang, F.X. (2018). RNA-Seq Analysis of Islets to Characterise the Dedifferentiation in Type 2 Diabetes Model Mice db/db. Endocr Pathol 29, 207–221. 10.1007/s12022-018-9523-x.

40. Egozi, A., Bahar Halpern, K., Farack, L., Rotem, H., and Itzkovitz, S. (2020). Zonation of Pancreatic Acinar Cells in Diabetic Mice. Cell Rep 32, 108043. 10.1016/j.celrep.2020.108043.

41. Jiang, Z., Wu, F., Laise, P., Takayuki, T., Na, F., Kim, W., Kobayashi, H., Chang, W., Takahashi, R., Valenti, G., et al. (2023). Tff2 defines transit-amplifying pancreatic acinar progenitors that lack regenerative potential and are protective against Kras-driven carcinogenesis. Cell Stem Cell 30, 1091–1109.e1097. 10.1016/j.stem.2023.07.002.

42. Tosti, L., Hang, Y., Debnath, O., Tiesmeyer, S., Trefzer, T., Steiger, K., Ten, F.W., Lukassen, S., Ballke, S., Kühl, A.A., et al. (2021). Single-Nucleus and In Situ RNA-Sequencing Reveal Cell Topographies in the Human Pancreas. Gastroenterology 160, 1330–1344.e1311. 10.1053/j.gastro.2020.11.010.

43. Chen, Z., Downing, S., and Tzanakakis, E.S. (2019). Four Decades After the Discovery of Regenerating Islet-Derived (Reg) Proteins: Current Understanding and Challenges. Front Cell Dev Biol 7, 235. 10.3389/fcell.2019.00235.

44. Li, Q., Wang, H., Zogopoulos, G., Shao, Q., Dong, K., Lv, F., Nwilati, K., Gui, X.Y., Cuggia, A., Liu, J.L., and Gao, Z.H. (2016). Reg proteins promote acinar-to-ductal metaplasia and act as novel diagnostic and prognostic markers in pancreatic ductal adenocarcinoma. Oncotarget 7, 77838–77853. 10.18632/oncotarget.12834.

45. Zhang, H., Corredor, A.L.G., Messina-Pacheco, J., Li, Q., Zogopoulos, G., Kaddour, N., Wang, Y., Shi, B.Y., Gregorieff, A., Liu, J.L., and Gao, Z.H. (2021). REG3A/REG3B promotes acinar to ductal metaplasia through binding to EXTL3 and activating the RAS-RAF-MEK-ERK signaling pathway. Commun Biol 4, 688. 10.1038/s42003-021-02193-z.

46. Cox, A.R., Lam, C.J., Rankin, M.M., King, K.A., Chen, P., Martinez, R., Li, C., and Kushner, J.A. (2016). Extreme obesity induces massive beta cell expansion in mice through self-renewal and does not alter the beta cell lineage. Diabetologia 59, 1231–1241. 10.1007/s00125-016-3922-7.

47. Linnemann, A.K., Baan, M., and Davis, D.B. (2014). Pancreatic beta-cell proliferation in obesity. Adv Nutr 5, 278–288. 10.3945/an.113.005488.

48. Zhao, H., Huang, X., Liu, Z., Pu, W., Lv, Z., He, L., Li, Y., Zhou, Q., Lui, K.O., and Zhou, B. (2021). Pre-existing beta cells but not progenitors contribute to new beta cells in the adult pancreas. Nat Metab 3, 352–365. 10.1038/s42255-021-00364-0.

49. Thorel, F., Damond, N., Chera, S., Wiederkehr, A., Thorens, B., Meda, P., Wollheim, C.B., and Herrera, P.L. (2011). Normal glucagon signaling and beta-cell function after near-total alpha-cell ablation in adult mice. Diabetes 60, 2872–2882. 10.2337/db11-0876.

50. Thorel, F., Nepote, V., Avril, I., Kohno, K., Desgraz, R., Chera, S., and Herrera, P.L. (2010). Conversion of adult pancreatic alpha-cells to beta-cells after extreme beta-cell loss. Nature 464, 1149–1154. 10.1038/nature08894.

51. Bonner-Weir, S., Inada, A., Yatoh, S., Li, W.C., Aye, T., Toschi, E., and Sharma, A. (2008). Transdifferentiation of pancreatic ductal cells to endocrine beta-cells. Biochem Soc Trans 36, 353–356. 10.1042/BST0360353.

52. Inada, A., Nienaber, C., Katsuta, H., Fujitani, Y., Levine, J., Morita, R., Sharma, A., and Bonner-Weir, S. (2008). Carbonic anhydrase II-positive pancreatic cells are progenitors for both endocrine and exocrine pancreas after birth. Proc Natl Acad Sci U S A 105, 19915–19919. 10.1073/pnas.0805803105.

53. Ackermann, A.M., Zhang, J., Heller, A., Briker, A., and Kaestner, K.H. (2017). High-fidelity Glucagon-CreER mouse line generated by CRISPR-Cas9 assisted gene targeting. Mol Metab 6, 236–244. 10.1016/j.molmet.2017.01.003.

54. Wicksteed, B., Brissova, M., Yan, W., Opland, D.M., Plank, J.L., Reinert, R.B., Dickson, L.M., Tamarina, N.A., Philipson, L.H., Shostak, A., et al. (2010). Conditional gene targeting in mouse pancreatic ß-Cells: analysis of ectopic Cre transgene expression in the brain. Diabetes 59, 3090–3098. 10.2337/db10-0624.

55. Madisen, L., Zwingman, T.A., Sunkin, S.M., Oh, S.W., Zariwala, H.A., Gu, H., Ng, L.L., Palmiter, R.D., Hawrylycz, M.J., Jones, A.R., et al. (2010). A robust and high-throughput Cre reporting and characterization system for the whole mouse brain. Nat Neurosci 13, 133–140. 10.1038/nn.2467.

56. Sachs, S., Bastidas-Ponce, A., Tritschler, S., Bakhti, M., Bottcher, A., Sanchez-Garrido, M.A., Tarquis-Medina, M., Kleinert, M., Fischer, K., Jall, S., et al. (2020). Targeted pharmacological therapy restores beta-cell function for diabetes remission. Nat Metab 2, 192–209. 10.1038/s42255-020-0171-3.

57. Carrano, A.C., Mulas, F., Zeng, C., and Sander, M. (2017). Interrogating islets in health and disease with single-cell technologies. Mol Metab 6, 991–1001. 10.1016/j.molmet.2017.04.012.

58. Chiou, J., Zeng, C., Cheng, Z., Han, J.Y., Schlichting, M., Miller, M., Mendez, R., Huang, S., Wang, J., Sui, Y., et al. (2021). Single-cell chromatin accessibility identifies pancreatic islet cell type- and state-specific regulatory programs of diabetes risk. Nat Genet 53, 455–466. 10.1038/s41588-021-00823-0.

59. Chen, C.W., Guan, B.J., Alzahrani, M.R., Gao, Z., Gao, L., Bracey, S., Wu, J., Mbow, C.A., Jobava, R., Haataja, L., et al. (2022). Adaptation to chronic ER stress enforces pancreatic beta-cell plasticity. Nat Commun 13, 4621. 10.1038/s41467-022-32425-7.

60. Salinno, C., Buttner, M., Cota, P., Tritschler, S., Tarquis-Medina, M., Bastidas-Ponce, A., Scheibner, K., Burtscher, I., Bottcher, A., Theis, F.J., et al. (2021). CD81 marks immature and dedifferentiated pancreatic beta-cells. Mol Metab 49, 101188. 10.1016/j.molmet.2021.101188.

61. Cheng, C.W., Villani, V., Buono, R., Wei, M., Kumar, S., Yilmaz, O.H., Cohen, P., Sneddon, J.B., Perin, L., and Longo, V.D. (2017). Fasting-Mimicking Diet Promotes Ngn3-Driven beta-Cell Regeneration to Reverse Diabetes. Cell 168, 775–788 e712. 10.1016/j.cell.2017.01.040.

62. Van de Casteele, M., Leuckx, G., Baeyens, L., Cai, Y., Yuchi, Y., Coppens, V., De Groef, S., Eriksson, M., Svensson, C., Ahlgren, U., et al. (2013). Neurogenin 3+ cells contribute to beta-cell neogenesis and proliferation in injured adult mouse pancreas. Cell Death Dis 4, e523. 10.1038/cddis.2013.52.

63. Rukstalis, J.M., and Habener, J.F. (2009). Neurogenin3: a master regulator of pancreatic islet differentiation and regeneration. Islets 1, 177–184. 10.4161/isl.1.3.9877.

64. Li, H.J., Kapoor, A., Giel-Moloney, M., Rindi, G., and Leiter, A.B. (2012). Notch signaling differentially regulates the cell fate of early endocrine precursor cells and their maturing descendants in the mouse pancreas and intestine. Dev Biol 371, 156–169. 10.1016/j.ydbio.2012.08.023.

65. Fang, Z., Liu, X., and Peltz, G. (2023). GSEApy: a comprehensive package for performing gene set enrichment analysis in Python. Bioinformatics 39 10.1093/bioinformatics/btac757.

66. Chen, E.Y., Tan, C.M., Kou, Y., Duan, Q., Wang, Z., Meirelles, G.V., Clark, N.R., and Ma’ayan, A. (2013). Enrichr: interactive and collaborative HTML5 gene list enrichment analysis tool. BMC Bioinformatics 14, 128. 10.1186/1471-2105-14-128.

67. Maestas, M.M., Ishahak, M., Augsornworawat, P., Veronese-Paniagua, D.A., Maxwell, K.G., Velazco-Cruz, L., Marquez, E., Sun, J., Shunkarova, M., Gale, S.E., et al. (2024). Identification of unique cell type responses in pancreatic islets to stress. Nat Commun 15, 5567. 10.1038/s41467-024-49724-w.

68. Lenzen, S. (2008). The mechanisms of alloxan- and streptozotocin-induced diabetes. Diabetologia 51, 216–226. 10.1007/s00125-007-0886-7.

69. Ahn, C., An, B.S., and Jeung, E.B. (2015). Streptozotocin induces endoplasmic reticulum stress and apoptosis via disruption of calcium homeostasis in mouse pancreas. Mol Cell Endocrinol 412, 302–308. 10.1016/j.mce.2015.05.017.

70. Xu, G., Chen, J., Jo, S., Grayson, T.B., Ramanadham, S., Koizumi, A., Germain-Lee, E.L., Lee, S.J., and Shalev, A. (2022). Deletion of Gdf15 Reduces ER Stress-induced Beta-cell Apoptosis and Diabetes. Endocrinology 163. 10.1210/endocr/bqac030.

71. Zhang, R., Kim, J.S., Kang, K.A., Piao, M.J., Kim, K.C., and Hyun, J.W. (2011). Protective Mechanism of KIOM-4 in Streptozotocin-Induced Pancreatic β-Cells Damage Is Involved in the Inhibition of Endoplasmic Reticulum Stress. Evid Based Complement Alternat Med 2011. 10.1155/2011/231938.

72. Han, H., Cho, J.W., Lee, S., Yun, A., Kim, H., Bae, D., Yang, S., Kim, C.Y., Lee, M., Kim, E., et al. (2018). TRRUST v2: an expanded reference database of human and mouse transcriptional regulatory interactions. Nucleic Acids Res 46, D380–d386. 10.1093/nar/gkx1013.

73. Urano, F., Wang, X., Bertolotti, A., Zhang, Y., Chung, P., Harding, H.P., and Ron, D. (2000). Coupling of stress in the ER to activation of JNK protein kinases by transmembrane protein kinase IRE1. Science 287, 664–666. 10.1126/science.287.5453.664.

74. Kaneto, H., Matsuoka, T.A., Nakatani, Y., Kawamori, D., Matsuhisa, M., and Yamasaki, Y. (2005). Oxidative stress and the JNK pathway in diabetes. Curr Diabetes Rev 1, 65–72. 10.2174/1573399052952613.

75. Consortium, E.P. (2012). An integrated encyclopedia of DNA elements in the human genome. Nature 489, 57–74. 10.1038/nature11247.

76. Luo, Y., Hitz, B.C., Gabdank, I., Hilton, J.A., Kagda, M.S., Lam, B., Myers, Z., Sud, P., Jou, J., Lin, K., et al. (2020). New developments on the Encyclopedia of DNA Elements (ENCODE) data portal. Nucleic Acids Res 48, D882–D889. 10.1093/nar/gkz1062.

77. Miguel-Escalada, I., Bonàs-Guarch, S., Cebola, I., Ponsa-Cobas, J., Mendieta-Esteban, J., Atla, G., Javierre, B.M., Rolando, D.M.Y., Farabella, I., Morgan, C.C., et al. (2019). Human pancreatic islet three-dimensional chromatin architecture provides insights into the genetics of type 2 diabetes. Nat Genet 51, 1137–1148. 10.1038/s41588-019-0457-0.

78. Greenwald, W.W., Chiou, J., Yan, J., Qiu, Y., Dai, N., Wang, A., Nariai, N., Aylward, A., Han, J.Y., Kadakia, N., et al. (2019). Pancreatic islet chromatin accessibility and conformation reveals distal enhancer networks of type 2 diabetes risk. Nat Commun 10, 2078. 10.1038/s41467-019-09975-4.

79. Sherman, M.H., and Beatty, G.L. (2023). Tumor Microenvironment in Pancreatic Cancer Pathogenesis and Therapeutic Resistance. Annu Rev Pathol 18, 123–148. 10.1146/annurev-pathmechdis-031621-024600.

80. Bell, R.H., Jr., and Strayer, D.S. (1983). Streptozotocin prevents development of nitrosamine-induced pancreatic cancer in the Syrian hamster. J Surg Oncol 24, 258–262. 10.1002/jso.2930240404.

81. Zhang, A.M.Y., Xia, Y.H., Lin, J.S.H., Chu, K.H., Wang, W.C.K., Ruiter, T.J.J., Yang, J.C.C., Chen, N., Chhuor, J., Patil, S., et al. (2023). Hyperinsulinemia acts via acinar insulin receptors to initiate pancreatic cancer by increasing digestive enzyme production and inflammation. Cell Metab 35, 2119–2135 e2115. 10.1016/j.cmet.2023.10.003.

82. Bang, S., Chung, H.W., Park, S.W., Chung, J.B., Yun, M., Lee, J.D., and Song, S.Y. (2006). The clinical usefulness of 18-fluorodeoxyglucose positron emission tomography in the differential diagnosis, staging, and response evaluation after concurrent chemoradiotherapy for pancreatic cancer. J Clin Gastroenterol 40, 923–929. 10.1097/01.mcg.0000225672.68852.05.

83. Kerk, S.A., Papagiannakopoulos, T., Shah, Y.M., and Lyssiotis, C.A. (2021). Metabolic networks in mutant KRAS-driven tumours: tissue specificities and the microenvironment. Nat Rev Cancer 21, 510–525. 10.1038/s41568-021-00375-9.

84. Ying, H., Kimmelman, A.C., Lyssiotis, C.A., Hua, S., Chu, G.C., Fletcher-Sananikone, E., Locasale, J.W., Son, J., Zhang, H., Coloff, J.L., et al. (2012). Oncogenic Kras maintains pancreatic tumors through regulation of anabolic glucose metabolism. Cell 149, 656–670. 10.1016/j.cell.2012.01.058.

85. Kim, H.T., Desouza, A.H., Umhoefer, H., Han, J., Anzia, L., Sacotte, S.J., Williams, R.A., Blumer, J.T., Bartosiak, J.T., Fontaine, D.A., et al. (2022). Cholecystokinin attenuates beta-cell apoptosis in both mouse and human islets. Transl Res 243, 1–13. 10.1016/j.trsl.2021.10.005.

86. Lavine, J.A., Kibbe, C.R., Baan, M., Sirinvaravong, S., Umhoefer, H.M., Engler, K.A., Meske, L.M., Sacotte, K.A., Erhardt, D.P., and Davis, D.B. (2015). Cholecystokinin expression in the beta-cell leads to increased beta-cell area in aged mice and protects from streptozotocin-induced diabetes and apoptosis. Am J Physiol Endocrinol Metab 309, E819–828. 10.1152/ajpendo.00159.2015.

87. Lavine, J.A., Raess, P.W., Stapleton, D.S., Rabaglia, M.E., Suhonen, J.I., Schueler, K.L., Koltes, J.E., Dawson, J.A., Yandell, B.S., Samuelson, L.C., et al. (2010). Cholecystokinin is up-regulated in obese mouse islets and expands beta-cell mass by increasing beta-cell survival. Endocrinology 151, 3577–3588. 10.1210/en.2010-0233.

88. Linnemann, A.K., Neuman, J.C., Battiola, T.J., Wisinski, J.A., Kimple, M.E., and Davis, D.B. (2015). Glucagon-Like Peptide-1 Regulates Cholecystokinin Production in beta-Cells to Protect From Apoptosis. Mol Endocrinol 29, 978–987. 10.1210/me.2015-1030.

89. Del Poggetto, E., Ho, I.L., Balestrieri, C., Yen, E.Y., Zhang, S., Citron, F., Shah, R., Corti, D., Diaferia, G.R., Li, C.Y., et al. (2021). Epithelial memory of inflammation limits tissue damage while promoting pancreatic tumorigenesis. Science 373, eabj0486. 10.1126/science.abj0486.

90. Alonso-Curbelo, D., Ho, Y.J., Burdziak, C., Maag, J.L.V., Morris, J.P.t., Chandwani, R., Chen, H.A., Tsanov, K.M., Barriga, F.M., Luan, W., et al. (2021). A gene-environment-induced epigenetic program initiates tumorigenesis. Nature 590, 642–648. 10.1038/s41586-020-03147-x.

91. Badgley, M.A., Kremer, D.M., Maurer, H.C., DelGiorno, K.E., Lee, H.J., Purohit, V., Sagalovskiy, I.R., Ma, A., Kapilian, J., Firl, C.E.M., et al. (2020). Cysteine depletion induces pancreatic tumor ferroptosis in mice. Science 368, 85–89. 10.1126/science.aaw9872.

92. Segerstolpe, A., Palasantza, A., Eliasson, P., Andersson, E.M., Andreasson, A.C., Sun, X., Picelli, S., Sabirsh, A., Clausen, M., Bjursell, M.K., et al. (2016). Single-Cell Transcriptome Profiling of Human Pancreatic Islets in Health and Type 2 Diabetes. Cell Metab 24, 593–607. 10.1016/j.cmet.2016.08.020.

93. Lawlor, N., George, J., Bolisetty, M., Kursawe, R., Sun, L., Sivakamasundari, V., Kycia, I., Robson, P., and Stitzel, M.L. (2017). Single-cell transcriptomes identify human islet cell signatures and reveal cell-type-specific expression changes in type 2 diabetes. Genome Res 27, 208–222. 10.1101/gr.212720.116.

94. Wang, Y.J., Schug, J., Won, K.J., Liu, C., Naji, A., Avrahami, D., Golson, M.L., and Kaestner, K.H. (2016). Single-Cell Transcriptomics of the Human Endocrine Pancreas. Diabetes 65, 3028–3038. 10.2337/db16-0405.

95. Muraro, M.J., Dharmadhikari, G., Grun, D., Groen, N., Dielen, T., Jansen, E., van Gurp, L., Engelse, M.A., Carlotti, F., de Koning, E.J., and van Oudenaarden, A. (2016). A Single-Cell Transcriptome Atlas of the Human Pancreas. Cell Syst 3, 385–394 e383. 10.1016/j.cels.2016.09.002.

96. Baron, M., Veres, A., Wolock, S.L., Faust, A.L., Gaujoux, R., Vetere, A., Ryu, J.H., Wagner, B.K., Shen-Orr, S.S., Klein, A.M., et al. (2016). A Single-Cell Transcriptomic Map of the Human and Mouse Pancreas Reveals Inter- and Intra-cell Population Structure. Cell Syst 3, 346–360 e344. 10.1016/j.cels.2016.08.011.

97. van der Meulen, T., Mawla, A.M., DiGruccio, M.R., Adams, M.W., Nies, V., Dolleman, S., Liu, S., Ackermann, A.M., Caceres, E., Hunter, A.E., et al. (2017). Virgin Beta Cells Persist throughout Life at a Neogenic Niche within Pancreatic Islets. Cell Metab 25, 911–926 e916. 10.1016/j.cmet.2017.03.017.

98. Lee, S., Zhang, J., Saravanakumar, S., Flisher, M.F., Grimm, D.R., van der Meulen, T., and Huising, M.O. (2021). Virgin beta-Cells at the Neogenic Niche Proliferate Normally and Mature Slowly. Diabetes 70, 1070–1083. 10.2337/db20-0679.

99. Ying, W., Lee, Y.S., Dong, Y., Seidman, J.S., Yang, M., Isaac, R., Seo, J.B., Yang, B.H., Wollam, J., Riopel, M., et al. (2019). Expansion of Islet-Resident Macrophages Leads to Inflammation Affecting beta Cell Proliferation and Function in Obesity. Cell Metab 29, 457–474 e455. 10.1016/j.cmet.2018.12.003.

100. Kulkarni, A., Muralidharan, C., May, S.C., Tersey, S.A., and Mirmira, R.G. (2022). Inside the beta Cell: Molecular Stress Response Pathways in Diabetes Pathogenesis. Endocrinology 164. 10.1210/endocr/bqac184.

101. Talchai, C., Xuan, S., Lin, H.V., Sussel, L., and Accili, D. (2012). Pancreatic beta cell dedifferentiation as a mechanism of diabetic beta cell failure. Cell 150, 1223–1234. 10.1016/j.cell.2012.07.029.

102. Wang, Z., York, N.W., Nichols, C.G., and Remedi, M.S. (2014). Pancreatic beta cell dedifferentiation in diabetes and redifferentiation following insulin therapy. Cell Metab 19, 872–882. 10.1016/j.cmet.2014.03.010.

103. Tenenbaum, M., Plaisance, V., Boutry, R., Pawlowski, V., Jacovetti, C., Sanchez-Parra, C., Ezanno, H., Bourry, J., Beeler, N., Pasquetti, G., et al. (2021). The Map3k12 (Dlk)/JNK3 signaling pathway is required for pancreatic beta-cell proliferation during postnatal development. Cell Mol Life Sci 78, 287–298. 10.1007/s00018-020-03499-7.

104. Dybala, M.P., Kuznetsov, A., Motobu, M., Hendren-Santiago, B.K., Philipson, L.H., Chervonsky, A.V., and Hara, M. (2020). Integrated Pancreatic Blood Flow: Bidirectional Microcirculation Between Endocrine and Exocrine Pancreas. Diabetes 69, 1439–1450. 10.2337/db19-1034.

105. Rupnik, M.S., and Hara, M. (2024). Local dialogues between the endocrine and exocrine cells in the pancreas. Diabetes. 10.2337/db23-0760.

106. Moon, K.R., van Dijk, D., Wang, Z., Gigante, S., Burkhardt, D.B., Chen, W.S., Yim, K., Elzen, A.v.d., Hirn, M.J., Coifman, R.R., et al. (2019). Visualizing structure and transitions in high-dimensional biological data. Nature Biotechnology 37, 1482–1492. 10.1038/s41587-019-0336-3.

107. Wolf, F.A., Angerer, P., and Theis, F.J. (2018). SCANPY: large-scale single-cell gene expression data analysis. Genome Biol 19, 15. 10.1186/s13059-017-1382-0.

108. van Dijk, D., Sharma, R., Nainys, J., Yim, K., Kathail, P., Carr, A.J., Burdziak, C., Moon, K.R., Chaffer, C.L., Pattabiraman, D., et al. (2018). Recovering Gene Interactions from Single-Cell Data Using Data Diffusion. Cell 174, 716–729 e727. 10.1016/j.cell.2018.05.061.

109. Luecken, M.D., Büttner, M., Chaichoompu, K., Danese, A., Interlandi, M., Mueller, M.F., Strobl, D.C., Zappia, L., Dugas, M., Colomé-Tatché, M., and Theis, F.J. (2022). Benchmarking atlas-level data integration in single-cell genomics. Nature Methods 19, 41–50. 10.1038/s41592-021-01336-8.

110. Zhang, Z., Mathew, D., Lim, T., Mason, K., Martinez, C.M., Huang, S., Wherry, E.J., Susztak, K., Minn, A.J., Ma, Z., and Zhang, N.R. (2023). Signal recovery in single cell batch integration. bioRxiv. 10.1101/2023.05.05.539614.

111. Venkat, A., Bhaskar, D., and Krishnaswamy, S. (2023). Multiscale geometric and topological analyses for characterizing and predicting immune responses from single cell data. Trends Immunol 44, 551–563. 10.1016/j.it.2023.05.003.

112. Li, D., Hsu, S., Purushotham, D., Sears, R.L., and Wang, T. (2019). WashU Epigenome Browser update 2019. Nucleic Acids Res 47, W158–W165. 10.1093/nar/gkz348.

113. Li, D., Purushotham, D., Harrison, J.K., Hsu, S., Zhuo, X., Fan, C., Liu, S., Xu, V., Chen, S., Xu, J., et al. (2022). WashU Epigenome Browser update 2022. Nucleic Acids Res 50, W774–W781. 10.1093/nar/gkac238.

114. Li, D., Harrison, J.K., Purushotham, D., and Wang, T. (2022). Exploring genomic data coupled with 3D chromatin structures using the WashU Epigenome Browser. Nat Methods 19, 909–910. 10.1038/s41592-022-01550-y.

115. Zhuo, X., Hsu, S., Purushotham, D., Kuntala, P.K., Harrison, J.K., Du, A.Y., Chen, S., Li, D., and Wang, T. (2023). Comparing genomic and epigenomic features across species using the WashU Comparative Epigenome Browser. Genome Res 33, 824–835. 10.1101/gr.277550.122.

